# Alveolar niche disruption and aberrant epithelial reprogramming are early hallmarks of idiopathic pulmonary fibrosis

**DOI:** 10.64898/2026.05.27.727792

**Authors:** Aurelien Justet, Venerino Poletti, Cristian Coarfa, Nebal Abu Hussein, Taylor Adams, Alan Waich, Agshin Balayev, Xiting Yan, Zhongyu Cai, Farah Moussa, Ruben De Man, Johad Khoury, Jonas C Schupp, Juan D Zuluaga, Amy Zhao, Julian Villalba, Farida Ahangari, Scott A Ochsner, Edward Manning, Wendy J Introne, Robert Homer, Bernadette R Gochuico, Laurens De Sadeleer, Claudia Carducci, Maria E Ruiz Echartea, Chao He, Bart M Vanaudenaerde, Wim Wuyts, Claudia Ravaglia, Ivan Rosas, Sara Tomassetti, Naftali Kaminski

**Author notes:** **Corresponding author:** Naftali Kaminski.

## Abstract

Idiopathic pulmonary fibrosis (IPF) is a progressive interstitial lung disease in which the earliest cellular events driving fibrosis remain poorly defined. Here, we analyzed lung samples from three independent and unique cohorts of patients with early disease and preserved lung function (Florence, NIH, Forli), applying an integrated multi-modal approach combining single-nucleus RNA sequencing, bulk transcriptomics, immunostaining, and spatial transcriptomics.

Single nuclear RNA sequencing of samples obtained by diagnostic bronchoscopic cryobiopsy (Florence, n= 22) revealed that early IPF is characterized by a marked shift in alveolar epithelial composition, with loss of AT1 and AT2 cells and the emergence of aberrant basaloid cells and alveolar epithelial intermediate cells. These populations exhibited transcriptional programs associated with epithelial plasticity and profibrotic signaling and closely resembled those observed in end-stage IPF. Higher proportions of aberrant basaloid and alveolar epithelial intermediate cells were associated with subsequent disease progression, whereas AT2 cell abundance correlated with preserved lung function. Fibrotic CTHRC1+ fibroblasts are largely restricted to advanced disease, while endothelial remodeling and inflammatory fibroblast states are already evident in early IPF. Spatial transcriptomic analyses confirmed early disruption of the alveolar niche, with replacement of normal epithelial–capillary interactions by aberrant epithelial and venous endothelial cells (Forli, n= 24); the findings were replicated through single cell RNA sequencing of samples obtained by video assisted thoracoscopy two decades earlier (NIH n=9).

Together, these findings identify that alveolar niche remodeling with loss of its normal components, and emergence of aberrant basaloid cells are features of early IPF, highlighting epithelial dysfunction as a key potential target for therapeutic interventions in early disease.

## Main

Progressive fibrosing interstitial lung diseases, including idiopathic pulmonary fibrosis (IPF), represent a major cause of respiratory morbidity and premature mortality worldwide ^1^. These disorders are characterized by relentless loss of lung function, progressive dyspnea, and mortality. Although antifibrotic therapies can attenuate the rate of decline, they do not halt disease progression or restore normal lung architecture, and their impact on long-term outcomes remains modest ^2–4^. The working model for IPF proposes multiple cycles of epithelial cell injury ^5^ in the context of aging ^6^ and genetic predisposition ^7,8^ that provoke the migration, proliferation ^9^, and activation of fibroblasts ^10^ and macrophages ^11^ and with the formation of active fibroblastic foci ^12^, accumulation of extracellular matrix (ECM), relentless lung remodeling and eventual respiratory failure. The application of single-cell and nucleus RNA sequencing (snRNA-seq) technologies have substantially enriched our understanding of the molecular and cellular underpinnings of human pulmonary fibrosis, uncovering the wide repertoire of changes in native cellular populations in the lung as well as previously unrecognized cellular populations and states ^10,13–21^, that were largely confirmed by subsequent spatial transcriptomic studies ^22,23^. However, these observations were mainly derived from samples obtained at end stage disease, as were previous bulk transcriptomic studies ^24,25^. This is of particular importance, as mounting evidence indicates that progressive pulmonary fibrosis develops well before clinical presentation. Within this framework, interstitial lung abnormalities (ILA), defined as incidental radiological findings on computed tomography in otherwise asymptomatic individuals^13,26^. While some ILAs may remain stable or reflect non-specific changes, others represent biologically active and progressive disease states. Clinical studies have begun to characterize these early stages by focusing on individuals at increased risk, particularly first-degree relatives of patients with familial pulmonary fibrosis (FPF) or familial interstitial pneumonia (FIP) ^27^. In these populations, early pulmonary fibrosis has been shown to be highly prevalent, with an incidence estimated to be more than 100-fold higher than that of sporadic idiopathic pulmonary fibrosis ^28^ and is associated with functional decline and reduced survival. In parallel, longitudinal imaging studies have demonstrated that even minimal interstitial abnormalities frequently represent subclinical disease with a high risk of progression. In relatives of patients with FPF, progression was observed in up to 75% of individuals with mild-to-moderate abnormalities ^29^. However, there is little information about the molecular and cellular characteristics of the lung in patients with early pulmonary fibrosis. To address this knowledge gap, we analyzed lung tissue obtained from three independent cohorts of patients with pulmonary fibrosis and preserved pulmonary functions. We performed single nuclear RNA sequencing (snRNASeq) on samples obtained by diagnostic bronchoscopic cryobiopsy from early disease patients at the University of Florence, and by Video-assisted thoracoscopic surgery (VATS) from asymptomatic relatives from Familial pulmonary fibrosis (FPF) at the NIH (Figure 1A), and confirmed the results by tissue staining, immunofluorescence and spatial transcriptomic analysis performed on a third cohort of samples obtained by diagnostic bronchoscopic cryobiopsy from early disease patients at the University of Forli. Our objective was to construct a detailed and temporally informed cellular atlas of the earliest steps of pulmonary fibrosis. By mapping the emergence, continuity, and diversification of cellular states in early disease, this approach aims to illuminate the sequence of biological events that precedes irreversible fibrosis, refine our understanding of how fibrotic niches are established, and provide a foundation for developing biomarkers and therapeutic strategies that target fibrosis at its inception rather than its end-stage manifestations.

**Figure 1:**
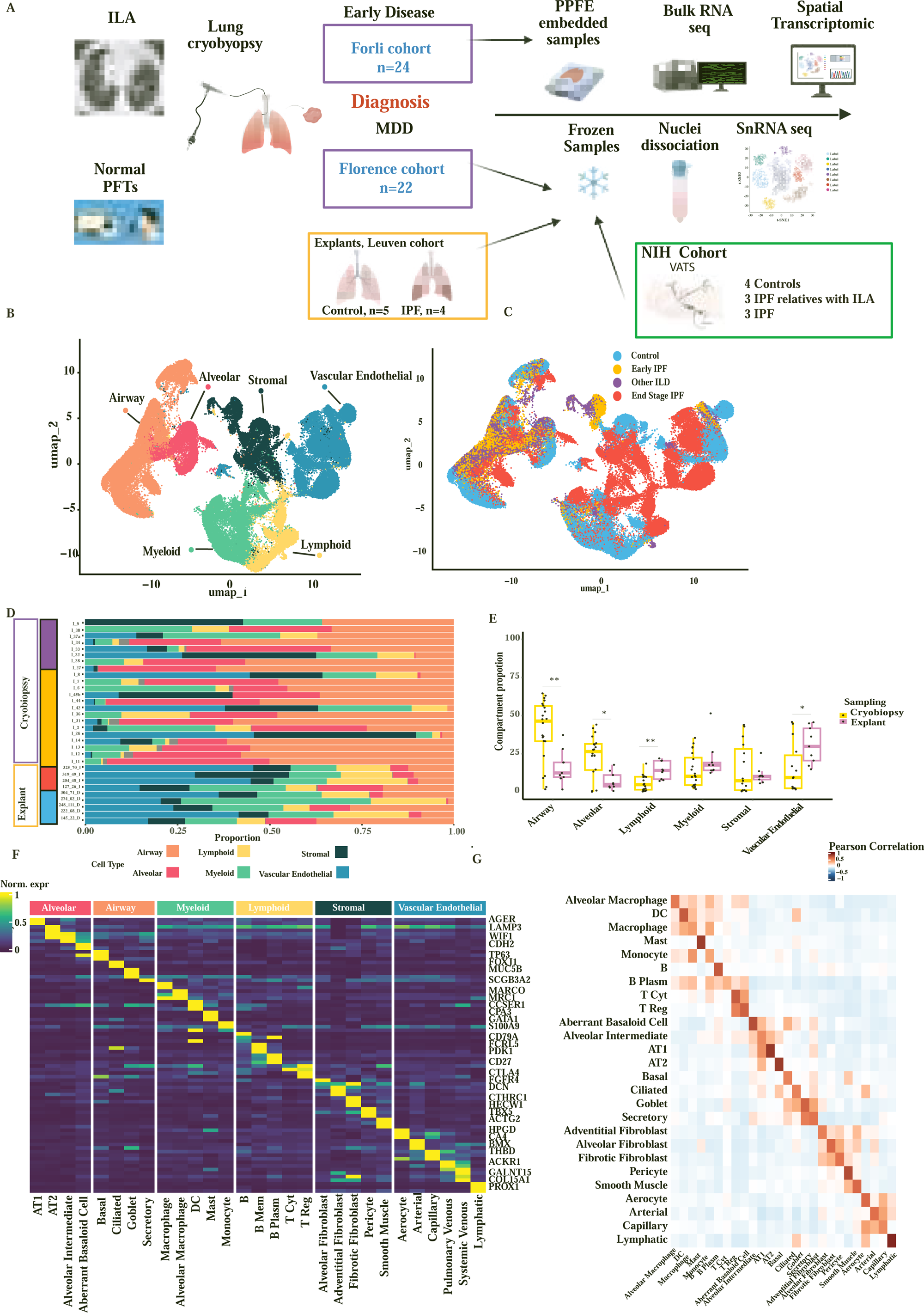
snRNA-seq of lung cryobiopsies reveals epithelial enrichment and preserved cellular diversity in early IPF. A) Overview of tissue procurement and experiment design. B) UMAPs of nuclei from integrated analysis of multiple datasets including Florence (Early ILDs) and Leuven cohorts (End Stage IPF and Controls), annotated per compartment. C) UMAPs of nuclei from integrated analysis of multiple datasets including Florence (Early ILDs) and Leuven cohorts (End Stage IPF and Controls), annotation based on the condition ie Early IPF, Others ILDs, End Stage IPF, D) Stacked boxplot E) Boxplots of cell type proportion normalized to the total number of cells per compartment F) Heat map of marker genes for all identified cell types, categorized into compartments. Each cell type is represented by the top three genes ranked by false discovery rate (FDR) adjusted P value. Gene expression values are unity normalized from 0 to 1 across rows within each categorical cell type group. G) Heatmap of spearman correlation between cell-type proportions.

## Results

We analyzed lung samples obtained by diagnostic bronchoscopic cryobiopsies from two independent Italian cohorts of patients with early interstitial lung disease defined by subnormal lung function test, recruited at the Careggi University Hospital, Florence (n = 22) and Department of Diseases of the Thorax, G.B. Morgagni Hospital/Bologna University Forlì (n = 24). The final diagnosis was established after multidisciplinary team discussion. Among these patients, 14 (63.6.%) and 12 (50%) were diagnosed with early IPF based on preserved FVC in the Florence and Forlì cohorts, respectively (Table 1). Other ILDs group included patients diagnosed with CTD-NSIP, Pleuro Parenchymal Fibro Elastosis, hypersensitivity pneumonitis, smoking related ILD and post COVID Fibrotic ILD, (Figure S1, A-F). The characteristics of the population are summarized in Table 1. Patients with early IPF were predominantly male, with a median age of 69.7 and 72.1 years in Florence and Forlì, respectively. Lung function tests were within the normal range at the time of diagnosis with no evidence of emphysema on CT scan. The mean follow-up was 2.0 years. No significant differences in baseline characteristics were observed between the two cohorts. Single-nucleus RNA-seq was performed on frozen samples from the Florence cohort, while bulk RNA-seq and spatial transcriptomics were conducted on paraffin-embedded samples from the Forlì cohort. To validate our findings, we leveraged an independent single-nucleus RNA sequencing (snRNA-seq) North American cohort of lung samples obtained by surgical (VATS) biopsies, established by the NIH ^27^. This cohort comprised relatives of patients with familial pulmonary fibrosis (FPF; n = 3) with preserved lung function, patients with sporadic idiopathic pulmonary fibrosis (IPF; n = 3), and healthy controls (n = 4).

**Table 1.** Characteristic of the population.

| Cohort | Forli |  |  | Florence |  |  | NIH |  |  |
| --- | --- | --- | --- | --- | --- | --- | --- | --- | --- |
|  | Over<br>all | Early<br>IPF | OtherILD | Over<br>all | Early<br>IPF | OtherILD | Ctl | Early<br>FPF | IPF |
| N, % | 24 | 12 | 12 | 22 | 14 | 8 | 4 | 3 | 3 |
| Age | 66.8 | 69.7 | 63.3 | 72±4 | 72 ±9 | 72±6 | 55±0 | 55.9±8 | 66±5 |
| % of<br>women | 42 | 29 | 54 | 41 | 33 | 55 | NA | 100 | 63 |
| FVC<br>(% of<br>pred) | 92+<br>-2 | 98<br>±13 | 84±<br>10 | 83±<br>16 | 88 ±<br>17 | 75±<br>10 | NA | NA | NA |
| DLCO (<br>% of<br>pred) | 85±<br>14 | 89.3±<br>10 | 82 ± 15 | 60±<br>12 | 64±<br>13 | 55+±<br>9 | NA | 74.6 | NA |

### Cryobiopsy SnRNA seq profiling uncovers heterogeneity of the fibrotic lung

SnRNA-seq of 24 frozen cryobiopsies from the Florence cohort, integrated with lung explants from 5 controls and 4 explants obtained from patients ontaining lung transplantation for end-stage IPF from the Leuven Cohort (GEO <u>GSE286182</u>), profiled 95542 nuclei (Figure 1B and C). We identified 4 different compartments including Airway Alveolar, Stromal, Myeloid, Lymphoid. We observed differences in compartment sampling between explant and cryobiopsy specimens, with relative depletion of vascular endothelial (p=0.02), and lymphoid populations, and enrichment of airway (p=0.006) and alveolar (p=0.01) epithelial populations in both early IPF and early ILD cryobiopsies (Figure 1D and E). There were no significant differences regarding myeloid (p=0.08) and stromal (p=0.53)

Despite the limited tissue availability from cryobiopsies, we identified all major cell types previously described in normal and fibrotic lungs, with strong correlation (mean R=0.62) to published Single Cell reference signatures ^30^ (Figure 1G and F).

### Early IPF is characterized by a shift of alveolar epithelial population and the emergence of aberrant basaloid cells

We analyzed 27199 epithelial nuclei and identified all major epithelial subtypes described in normal and fibrotic lungs ^31^, including aberrant basaloid cells (ABBA) and an alveolar intermediate state positioned between AT1, AT2, and ABBA (Figure 2A). As previously described ^13,32,33^, ABBA expressed basal markers (TP63, KRT1), senescence hallmarks (CDKN2A), TGFβ-inducible genes (EPHB2, CASC15, TGFBI), extracellular matrix (ECM) remodeling genes (COL1A1, COL5A1, PRSS2,), the epithelial–mesenchymal transition (EMT) hallmark CDH2 and a biomarker of IPF progression (MMP7). Alveolar intermediates displayed mixed molecular features of AT1 (GPC5, CLIC5), AT2 (WIF1, LRKK2), and basaloid cells (CTSE, CPA6, FRMD5) but lacked canonical basaloid markers (TP63), EMT markers, and senescence markers (CDKN1A, CDKN2A), (Figure 2B). While these cells were always found in End Stage IPF, Aberrant basaloid and alveolar intermediate cells were detected in 64.3% and 93% in patients with IPF, respectively (Figure 2C). Compared with control lungs, both early and end-stage fibrosis showed a reduction in alveolar epithelial type 2 (AT2) cells (Early IPF 50.4% 69.4%, p = 0.03; End Stage IPF 57.4% vs 69.4%, p= 0.06) and type 1 (Early IPF 16.9% 30.6%, p = 0.007; End Stage IPF 19.3% vs 30.6%, p= 0.02) associated with the emergence alveolar intermediates (Early IPF 30.0% vs 0.38%, p = 0.004; End Stage IPF 8.6% vs 0.38%, p= 0.03) and ABBA (Early IPF 4.4% vs 0.05 %, p = 0.04; End Stage IPF 14,4% vs 0.05%, p= 0.0005). There was a significant increase in alveolar intermediate proportion in Ealy IPF as compared to End Stage IPF (p=0.04). Immunofluorescent staining validated our transcriptional signatures and confirmed the different alveolar states’ existence and location. ABBA co-expressed AT1 (AGER) or AT2 (SPC) markers together with Cytokeratin 17 (CK17), while alveolar intermediates expressed AT1/AT2 markers along with Cathepsin E (CTSE) (Figure 2F) as previously described ^32^. We labelled cells positive for CK17 and CTSE, expression by immunostaining (Figure 3A). CK17 positive cells were only found in the airway in the control group with no CTSE positive cells. Double positive cells were only found in the alveolar space in patients with early and end stage disease, in a similar proportion. Quantitative analysis revealed that the proportion of Cytokeratin 17- and CTSE-positive cells was significantly higher in both early disease (p=0.02) and end-stage fibrosis (p= 0.03) compared with controls, confirming the early increasing number ABBA during lung fibrogenesis (Figure 3B). Transcriptomic profiles of epithelial cell types in early ILD showed a strong correlation with those from end-stage disease (Figure 3C), indicating that major disease-associated programs are already established at early stages. Despite this overall concordance, several key pathways differed in magnitude between stages. End stage disease showed a significant higher expression of epithelial–mesenchymal transition (EMT) and TGFβ-responsive genes, as well as signatures related to angiogenesis, coagulation, and extracellular matrix remodeling, whereas PI3K–AKT–mTOR signaling appeared more prominent in Early IPF (Figure 3D, E). We next examined whether epithelial composition was associated with clinical progression. Patients were stratified according to progression status based on ATS/ERS criteria. Individuals who experienced disease progression during follow-up exhibited higher proportions of ABBA (7.7% vs 1.76%, p = 0.03) and alveolar intermediate cells (25.7% vs 9.3, p=0.03) compared with stable patients (Figure 3F). We identified a similar but non-significant trend, with a correlation with the lung function decline and the time to progression with the proportion of ABBA. In contrast, AT2 cell abundance was negatively correlated with FVC decline suggesting that changes of alveolar epithelium contribute to functional deterioration (Figure 3G).

**Figure 2:**
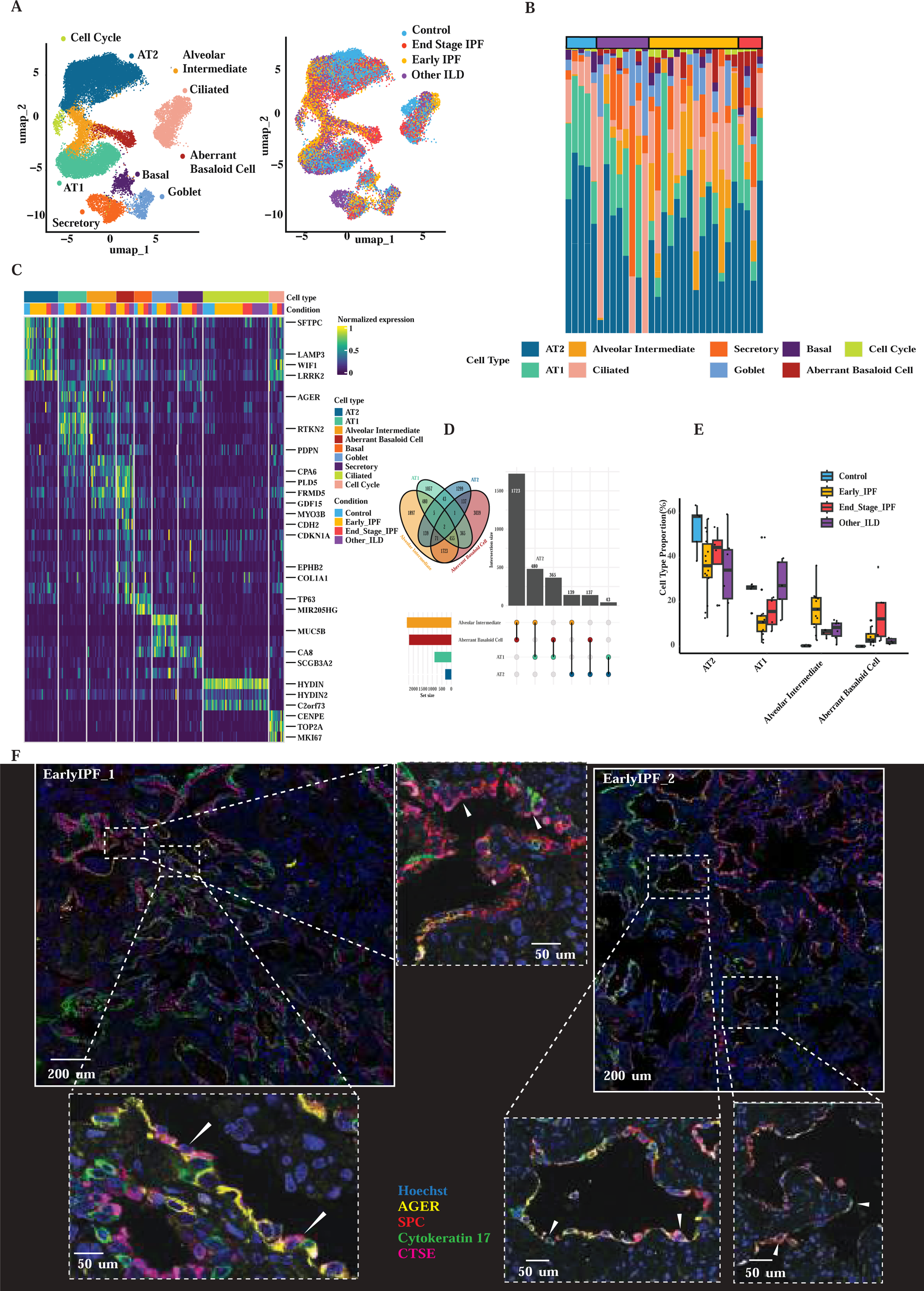
Early epithelial remodeling in IPF is marked by the emergence of alveolar intermediate and aberrant basaloid cells. A, UMAP representation of epithelial cell populations identified by single-cell transcriptomics, including AT1, AT2, alveolar intermediate, aberrant basaloid, basal, goblet, secretory, ciliated, and cycling epithelial cells. Right panel shows the distribution of cells colored by disease condition (control, early IPF, end-stage IPF and other ILD). B, Stacked bar plot showing the relative composition of epithelial cell populations across individual samples and disease conditions. C, Heat map of normalized expression of representative marker genes across epithelial cell populations, highlighting transcriptional programs distinguishing alveolar, airway and aberrant epithelial states. D, Overlap of differentially expressed genes among AT2, AT1, alveolar intermediate and aberrant basaloid epithelial populations (Venn diagram and UpSet plot), illustrating shared and cell-type–specific transcriptional signatures. E, Proportion of alveolar epithelial cell populations across disease groups, showing reduced AT1 and AT2 cells and increased alveolar intermediate and aberrant basaloid cells in IPF. Box plots indicate median, interquartile range and individual samples. F, Representative immunofluorescence images from early IPF lungs showing epithelial remodeling within alveolar regions. Staining for AGER (AT1), SPC (AT2), CK17 and CTSE highlights the presence of aberrant epithelial populations within remodeled alveolar structures. Insets show higher-magnification views. Scale bars, 200 μm and 50 μm

**Figure 3.**
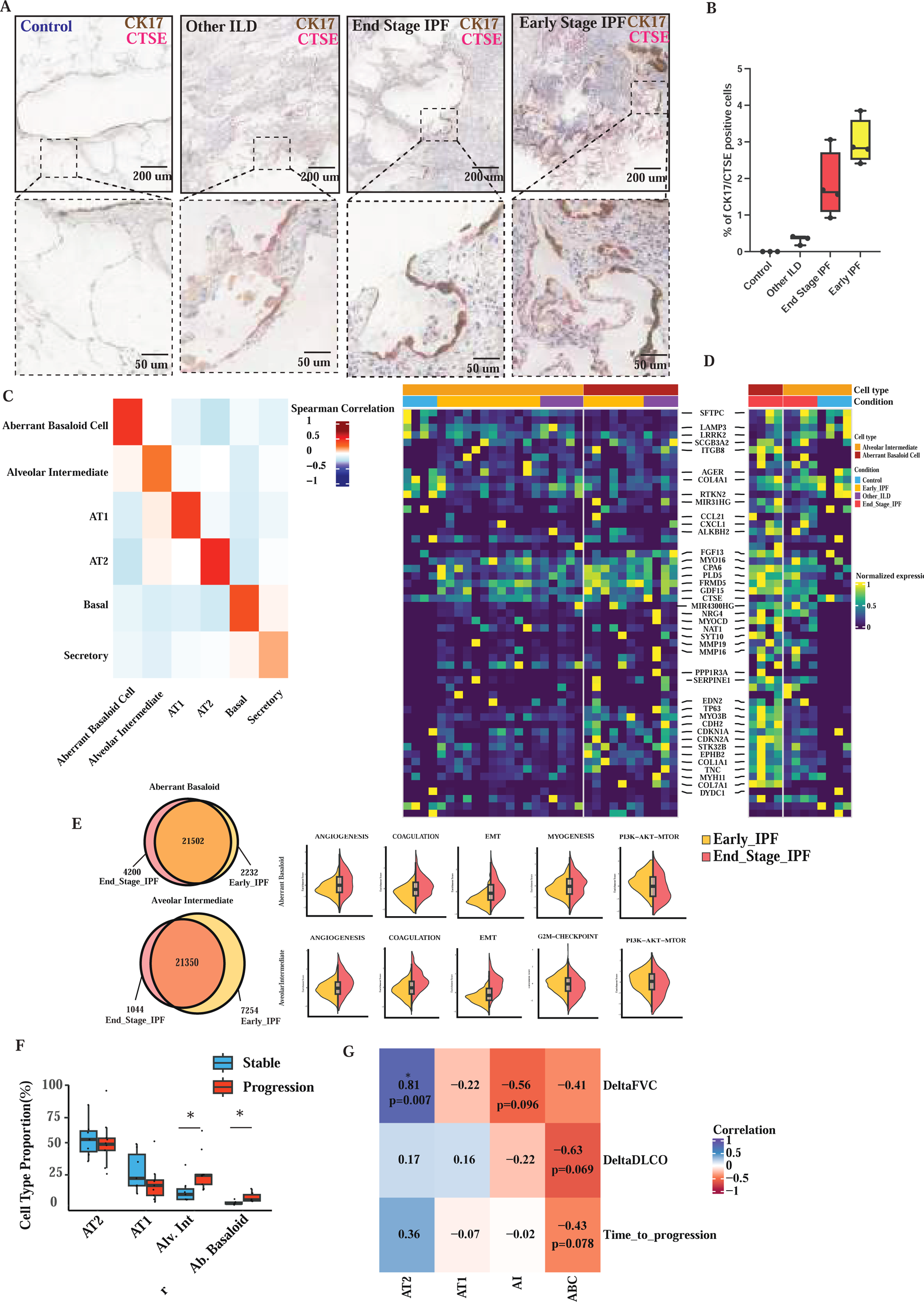
Emergence of aberrant basaloid and alveolar intermediate epithelial states in IPF and their association with disease progression. **A)** Representative immunohistochemistry for CK17 (brown) and CTSE (magenta) in lung sections from control, other ILD, end-stage IPF, and early-stage IPF lungs. Dashed boxes indicate regions shown at higher magnification. Scale bars, 200 μm (top) and 50 μm (bottom). **B**, Quantification of the proportion of CK17⁺CTSE⁺ epithelial cells across disease groups. Box plots indicate median, interquartile range and individual samples. **C**, Spearman correlation matrix showing transcriptional similarity between epithelial cell populations from Early and End Stage IPF. **D**, Heat map showing normalized expression of selected genes across epithelial cell types and disease conditions (control, early IPF, other ILD, end-stage IPF), highlighting distinct transcriptional programs between alveolar intermediate and aberrant basaloid cells. **E**, Overlap of differentially expressed genes between early IPF and end-stage IPF in aberrant basaloid and alveolar intermediate cells (Venn diagrams). Violin plots show enrichment scores for hallmark pathways including angiogenesis, coagulation, epithelial–mesenchymal transition, myogenesis, and PI3K–AKT–mTOR signaling. **F,** Proportion of epithelial cell types in patients with stable disease or disease progression, highlighting enrichment of alveolar intermediate and aberrant basaloid cells in progressive disease. Box plots indicate median, interquartile range and individual samples; asterisks denote statistical significance. **G**, Correlation between epithelial cell-type abundance and clinical outcomes (ΔFVC, ΔDLCO, and time to progression). AT2 cell proportion positively correlates with preservation of lung function, whereas alveolar intermediate and aberrant basaloid states tend to associate with worse clinical outcomes. Spearman correlation coefficients and *P* values are shown.

Together, these findings support a model in which early epithelial remodeling—marked by expansion of aberrant basaloid and transitional alveolar states and depletion of AT1 and AT2 cells—precedes overt fibrosis and is linked to subsequent disease progression. These epithelial states may therefore serve as early prognostic biomarkers and potential therapeutic targets aimed at preserving alveolar integrity.

### Early and end-stage IPF differ exhibit similar endothelial and immune but distinct mesenchymal changes

We analyzed 39180 nuclei from stromal and vascular endothelial populations (Figure 4A). The stromal compartment included smooth muscle cells, pericyte, alveolar and adventitial fibroblasts, as well as fibrotic fibroblasts, recently described and characterized by enrichment of COL1A1, COL3A1, CTHRC1 or POSTN (Figure 4B,4C). Fibrotic fibroblasts were detected ,predominantly, in end-stage IPF and in a patient with early Pleuroparenchymal Fibroelastosis, but were barely detected in early IPF cryobiopsies (Figure 4B, 4C). Notably, adventitial fibroblasts and smooth muscle cells were increased in early IPF and other ILD groups as compared to Control, likely reflecting enrichment of airway-associated cells in the sampled regions. Notably, both early and end-stage IPF displayed an increased proportion of alveolar fibroblasts compared with controls and other ILDs. While alveolar fibroblasts shared many commonly expressed genes across both conditions, alveolar fibroblasts from early IPF exhibited an enrichment of inflammation-associated genes, including SFRP4, CXCL12, and CXCL14, along with a reduction in profibrotic markers such as POSTN. Pathway analysis confirmed these observations, showing that alveolar fibroblasts from early IPF patients were enriched for inflammatory pathways (TNF alpha, IL6-JAK-STAT) and displayed a decrease in profibrotic pathways (TGF Beta, Wnt signaling) (Figure 4I).

**Figure 4.**
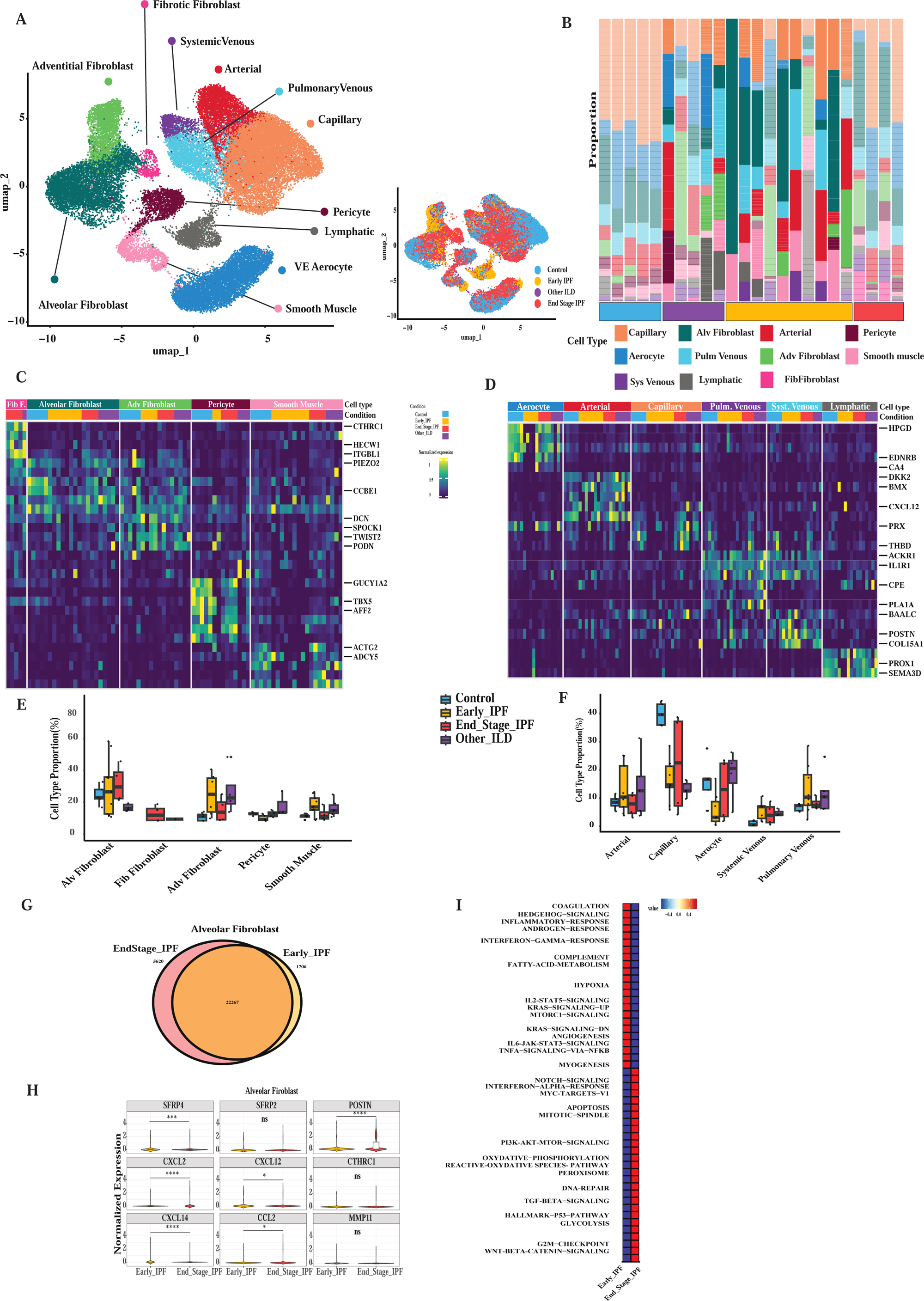
Remodeling of the immune microenvironment in IPF. **A**, UMAP representation of immune cell populations identified by single-cell transcriptomics, including alveolar macrophages, macrophages, monocytes, dendritic cells (DC), mast cells, B cells, plasma B cells, memory B cells, regulatory T cells (Treg) and cytotoxic T cells (T cyt). Bottom panel shows distribution of cells colored by disease condition (control, early IPF, other ILD, end-stage IPF). **B**, Heat map showing normalized expression of representative marker genes across immune cell populations and disease conditions, highlighting transcriptional signatures defining each immune subset. C, Relative proportions of innate immune populations (alveolar macrophages, macrophages, monocytes, dendritic cells and mast cells) across disease groups (control, early IPF, end-stage IPF and other ILD). Box plots indicate median, interquartile range and individual samples. D, Distribution of adaptive immune populations (B cells, plasma B cells, memory B cells, cytotoxic T cells and regulatory T cells) across disease conditions. **E**, Heat map showing transcriptional heterogeneity within alveolar macrophages, comparing control and early IPF samples and highlighting genes associated with inflammatory and remodeling programs. F Hallmark pathway enrichment analysis in alveolar macrophages, showing activation of inflammatory, metabolic and stress-related pathways including **T**NFα signaling via NF-κB, IL6–JAK–STAT3 signaling, oxidative phosphorylation and interferon responses. G, Proportion of pro-fibrotic alveolar macrophages compared with canonical alveolar macrophages in early IPF and end-stage IPF, highlighting expansion of the pro-fibrotic macrophage population during disease progression. ***P < 0.001

Within the vascular endothelial compartment, we identified aerocytes, general capillary endothelial cells, venous endothelial cells, arterial, and smooth muscle cells (Fig. 4a). Compared with controls, early IPF lungs showed a marked reduction in general capillary endothelial cells (14.0% vs. 40.0%, p = 0.003) and aerocytes (3.4% vs. 16.9%, p = 0.05), together with an expansion of systemic venous endothelial cells (0.5% vs. 7.7%, p = 0.01). This pattern was largely retained in end-stage IPF. Despite limitations related to the sampling of the immune compartment, snRNA-seq analysis revealed substantial heterogeneity within this compartment (Figure 5A, 5B). The proportion of alveolar macrophages was increased in both early and end-stage IPF compared with controls. In a subset of patients with early IPF, alveolar macrophages displayed a distinct profibrotic transcriptional signature characterized by increased expression of SPP1, CHI3L1, CHIT1 and CCL22 compared with controls (Figure 5E). Differential pathway analysis of alveolar macrophages from early versus end-stage IPF revealed enrichment of inflammatory and fibrotic programs in early disease (Figure 5F). Consistently, the proportion of alveolar macrophages co-expressing at least two of these markers was significantly higher in early IPF than in end-stage IPF (Figure 5G).

**Figure 5.**
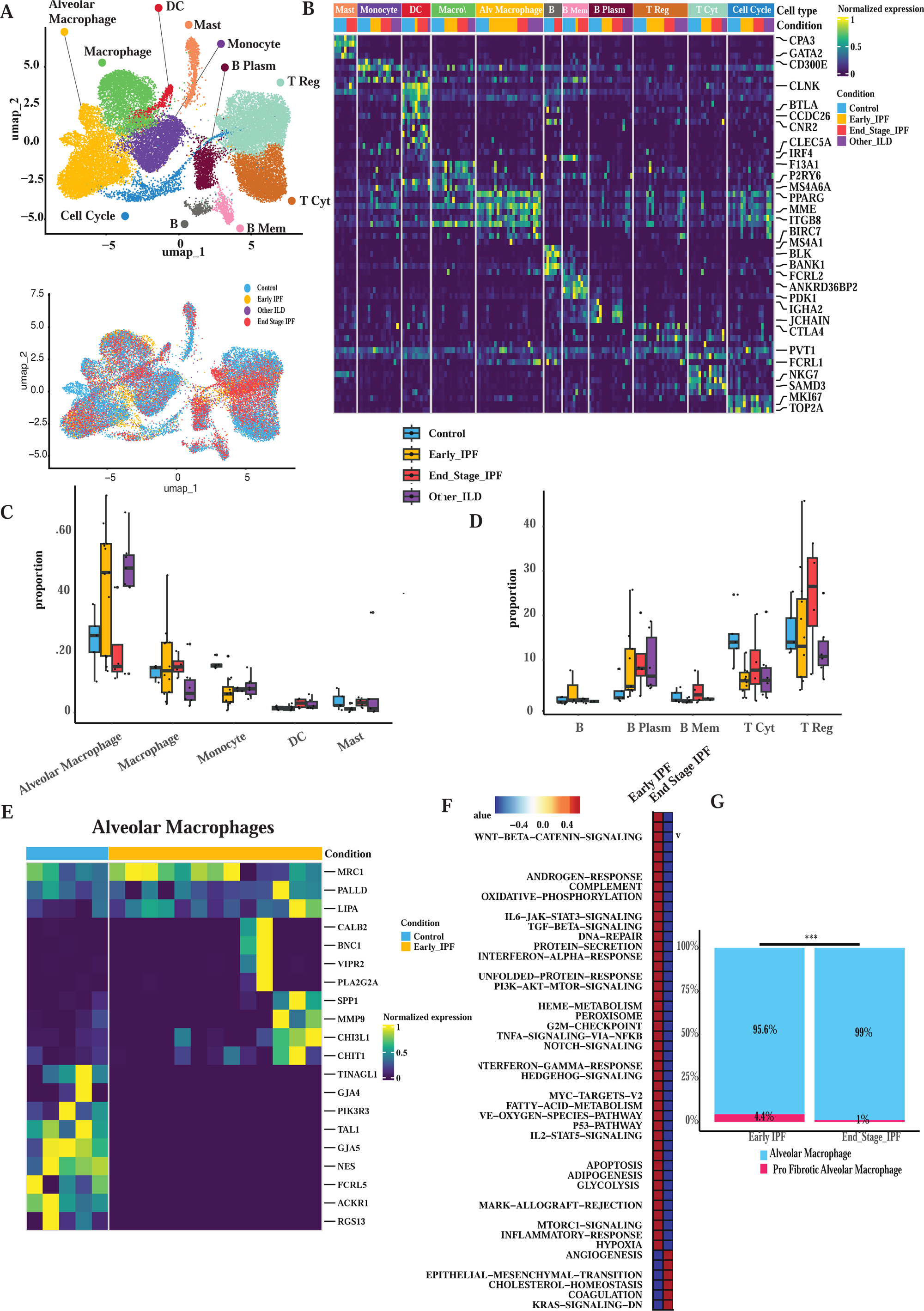
Stromal and vascular remodeling during IPF progression. **A**, UMAP representation of stromal and vascular cell populations identified by single-cell transcriptomics, including alveolar fibroblasts, fibrotic fibroblasts, adventitial fibroblasts, pericytes, smooth muscle cells, and endothelial populations (capillary, aerocyte, arterial, pulmonary venous, systemic venous and lymphatic endothelial cells). Bottom panel shows distribution of cells colored by disease condition. **B**, Stacked bar plot showing the relative composition of stromal and vascular cell populations across individual samples. **C**, Heat map of normalized expression of marker genes defining stromal cell populations, including alveolar fibroblasts, fibrotic fibroblasts, adventitial fibroblasts, pericytes and smooth muscle cells. **D**, Heat map showing gene expression signatures distinguishing endothelial subtypes (aerocyte, arterial, capillary, pulmonary venous, systemic venous and lymphatic endothelial cells) across disease conditions. **E**, Proportion of stromal cell populations across disease groups, highlighting expansion of fibrotic fibroblasts in IPF lungs. **F**, Relative abundance of endothelial cell populations across conditions. **G**, Venn diagram showing overlap of differentially expressed genes in alveolar fibroblas**ts** between early IPF and end-stage IPF. **H**, Differential expression of representative fibrosis-associated genes (SFRP1, POSTN, CCL2, CXCL12, CTHRC1, MMP11) in alveolar fibroblasts between early IPF and end-stage IPF. **I**, Hallmark pathway enrichment analysis in alveolar fibroblasts, highlighting activation of TGF-β signaling, IL6**–**JAK–STAT3 signaling, hypoxia, inflammatory response and extracellular matrix remodeling pathways during disease progression.

Together, these findings suggest the presence of temporally distinct remodeling patterns across immune, stromal and vascular endothelial compartments in IPF.

### Independent cohort analysis corroborates remodeling of the alveolar niche and the presence of aberrant basaloid cells as a feature of early pulmonary fibrosis

We next leveraged the NIH single-nucleus RNA sequencing dataset as an independent validation cohort to assess the robustness of our findings This dataset was generated from lung samples obtained by surgical (VATS) biopsies close to 20 years ago ^27^ and included asymptomatic relatives of patients with familial pulmonary fibrosis, patients with sporadic idiopathic pulmonary fibrosis, and healthy controls. Quality control metrics were comparable across cohorts, supporting the overall robustness of the dataset (Figure S1E). Unsupervised clustering resolved all major lung compartments, including epithelial, immune, stromal, and vascular populations, with consistent annotation supported by canonical marker genes (Figure S2 A-D). However, we observed a relatively limited representation of vascular endothelial, immune, and stromal populations, likely reflecting sampling bias (Figure S1D). We therefore focused subsequent analyses on epithelial populations, which enabled robust cross-cohort comparisons. Projection by condition revealed enrichment of specific cell populations in disease states, while correlation analyses confirmed transcriptional coherence within lineages (Figure 6 A-B). Importantly, direct quantification of cell-type proportions demonstrated reproducible shifts in cellular composition (Figure 6 C-D), Direct quantification of epithelial cell-type proportions revealed a decrease in AT1 cells, accompanied by an increased proportion of alveolar intermediate cells in relatives of patients with familial pulmonary fibrosis. Notably, ABBA were exclusively detected in pathological conditions (Figure 6E). Together, these findings support disease-associated epithelial remodeling in this independent cohort.

**Figure 6.**
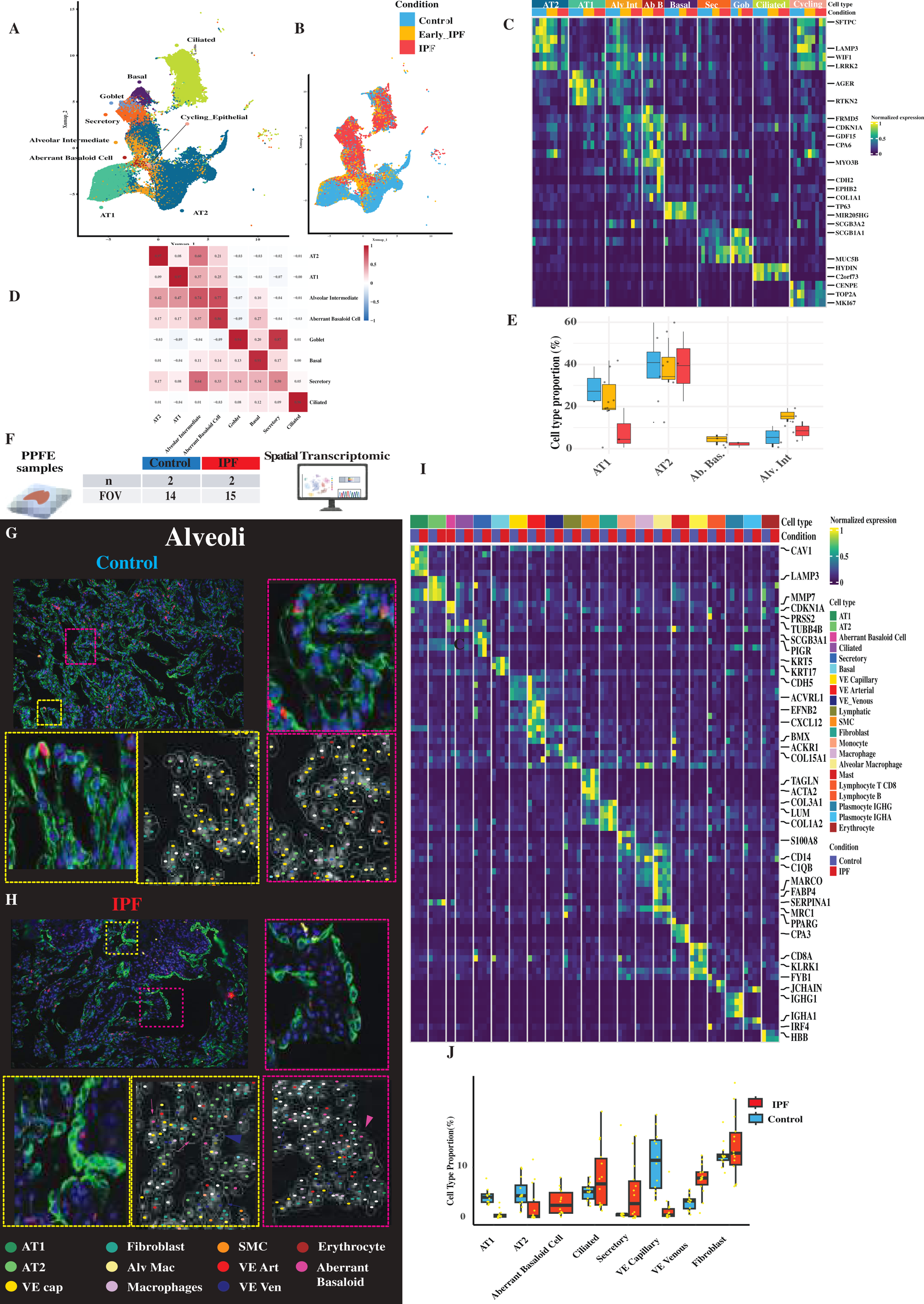
Cross-cohort validation of epithelial remodeling and spatial organization in pulmonary fibrosis. **(A)** UMAP representation of epithelial cells, identifying major epithelial populations including AT1, AT2, alveolar intermediate, aberrant basaloid, and airway epithelial subsets (basal, secretory, goblet, and ciliated cells).**(B)** UMAP colored by condition (Control, early FPF, and IPF), highlighting disease-associated shifts in epithelial cell states**.(C)** Correlation matrix of epithelial transcriptional profiles, revealing relationships between alveolar and airway epithelial populations**.(D)** Heatmap of selected marker genes displayed as Z-score–scaled expression, illustrating distinct transcriptional programs across epithelial cell types and confirming the presence of disease-associated states. **(E)** Cell-type proportions derived from single-nucleus RNA-seq data, showing relative abundance of epithelial populations across conditions and highlighting disease-associated changes, including expansion of intermediate and aberrant basaloid cells**.(F)** Spatial transcriptomics workflow and representative images of alveolar regions from control **(G)** and IPF lungs **(H).** Immunofluorescence images highlight cellular composition of alveolar niches, including AT1, AT2, vascular endothelial cells, fibroblasts, macrophages and smooth muscle cells. Insets show higher-magnification views and segmentation used for spatial cell identification**.(I),** Heat map showing normalized expression of representative marker genes across cell types identified in the spatial transcriptomic dataset, including epithelial, immune, stromal and vascular populations**.(J),** Quantification of the proportion of major cell populations within alveolar regions in control and IPF lungs, including AT1, AT2, aberrant basaloid cells, ciliated cells, secretory cells, endothelial cells and fibroblasts. Box plots indicate median, interquartile range and individual fields of view.

Bulk RNA-seq analysis of the Forlì cohort confirmed these observations, revealing enrichment significant enrichment of genes expressed by ABBA (ABC), including MMP7, GDF15, CTSE, CDH2, CDKN1A (Figure S3A. B, Table S1). To further refine these observations, we leveraged previously published single-cell datasets^32^ to guide interpretation. Deconvolution confirmed the trend observed in the Forli cohort including a decrease of alveolar epithelial cells and increased of ABBA (Figure S2C).

### Spatial transcriptomic highlight the change occurring in the alveolar Niche

Finally, we applied high-resolution probe-based spatial transcriptomic technology to two early IPF and two control samples, analyzing 14 and 15 fields of view, respectively (Figure 6F). This approach enabled the identification of the major cell types of the fibrotic lung (Figure 6 G-I) recapitulated the proximal and distal lung architecture (Figure 6C, Figure S4 A-C) and confirmed IFC results. Cell type proportion analysis across the different fields of view revealed a reduction of the alveolar epithelial population in early IPF compared with controls, the emergence of ABBA, and a decrease in general capillary cells accompanied by an expansion of venous endothelial cells (Figure 6J).

In control lungs, AT2 cells were primarily surrounded by AT1 cells, other AT2 cells, macrophages, and general capillaries, reflecting the expected organization of the healthy alveolar niche (Figure S4 C,D). In contrast, this spatial organization was profoundly altered in early IPF lungs, where AT2 cells were frequently neighbored by ABBA and venous endothelial cells, at the expense of AT1 cells and general capillaries. Delaunay network–based approach, which constructs a geometric graph linking each cell to its nearest neighbors, enabling systematic quantification of cellular neighborhoods and spatial relationships without prior assumptions revealed a marked reorganization of the alveolar niche in early IPF. Network-based quantification of the ten closest neighbors of AT2 cells confirmed these alterations, identifying venous endothelial cells as the third closest neighbor and ABBA as the ninth closest neighbor in early IPF lungs. These spatial associations were not observed in control samples. Subsequently, ligand–receptor interactions were inferred using our single-nucleus RNA-sequencing dataset. Putative signaling interactions were identified by examining the expression of ligands and the corresponding receptors in cell populations identified as components of the alveolar niche in early IPF. Within the spatially defined alveolar niche of the fibrotic lung, aberrant epithelial cells emerged as a major source of profibrotic cytokine signaling, whereas alveolar fibroblasts appeared to be the principal target cell population. In particular, abnormal epithelial populations actively engage in the secretion of signaling molecules strongly implicated in fibrogenesis. (TGFB, PDGF, WNT, CDH), whereas alveolar fibroblasts predominantly receive and integrate these signals and influence ECM production (Figure 6I).

Overall, spatial transcriptomic analysis corroborated the compositional changes observed by snRNA-seq, including reduced AT1 and AT2 cell representation and increased venous endothelial and aberrant basaloid cell populations. Together, these findings demonstrate that alveolar niche remodeling is an early event in IPF, preceding end-stage fibrosis and potentially contributing to disease initiation and progression.

## Discussion

In this study, we integrated cryobiopsy and VATS-based single-nucleus RNA sequencing, bulk transcriptomics, and high-resolution spatial transcriptomics across three independent patient cohorts to define the earliest cellular and tissue-level events in idiopathic pulmonary fibrosis (IPF). Our findings indicate that early IPF is reproducibly characterized primarily by profound loss of the epithelial and endothelia components of the alveolar niche, and the emergence of aberrant basaloid cells and intermediate alveolar epithelial cells.

A central observation of our work is the early and consistent emergence of aberrant basaloid cells and alveolar intermediate epithelial states, accompanied by a marked loss of AT2 and AT1 cells. While these populations have been previously described in end-stage IPF^13,16,19,20,23,32^, Our data demonstrate that these alterations are already present at early stages of the disease, when patients have preserved lung functions, without substantial accumulation of CTHRC1 fibroblasts and may be associated with disease progression. This suggests that maladaptive epithelial differentiation is not merely a consequence of fibrosis but instead may be a pathogenetic feature of the disease, contributing to its initiation and progression. The inverse correlation between AT2 abundance and lung function decline further supports a model in which loss of regenerative epithelial capacity is a key determinant of progression. These findings could have implications for current therapeutic strategies. Approved antifibrotic agents, including nintedanib ^2^,pirfenidone ^4^ ,nerandomilast ^3^ or trepoprostinil ^34^ primarily target fibroblast proliferation and extracellular matrix deposition. While these therapies slow disease progression, they do not restore alveolar architecture or halt disease onset. Our data suggests that interventions exclusively targeting fibroblasts may be intrinsically limited when applied early in the disease course. Instead, our results support a shift toward therapeutic strategies aimed at preserving epithelial integrity and restoring niche homeostasis. The early expansion of aberrant epithelial states and their association with progression further raise the possibility that these populations could be used to identify biomarkers for risk stratification and early intervention. Of interest, MMP7, a validated outcome predictive peripheral blood biomarker in idiopathic pulmonary fibrosis and progressive fibrotic lung disease^35^, is mainly expressed by aberrant basaloid cells in the lung and can also be observed in early interstitial lung disease ^36^. Our results would suggest assessing the value of MMP7 as well as other aberrant basaloid cell and intermediate epithelial cell markers such as GDF15, to determine the risk of progression in patients with early pulmonary fibrosis.

Beyond transcriptional changes, our spatial analyses reveal that early IPF is characterized by a profound reorganization of the alveolar niche. In healthy lungs, AT2 cells reside within a structured microenvironment supported by AT1 cells, general capillary endothelial cells, and resident macrophages. In early IPF, this organization is disrupted, with aberrant basaloid cells and venous endothelial cells replacing general capillaries in the immediate AT2 neighborhood. Such spatial remodeling is likely to have important functional consequences, including altered angiocrine signaling, impaired oxygen and nutrient exchange, and sustained epithelial stress. Notably, similar alterations have been described in end-stage IPF and in other fibrotic or inflammatory interstitial lung diseases ^33,37,38^; however, our findings indicate that these processes are already established at early stages of disease.

Several limitations should be acknowledged. The analysis of early human lung disease is inherently constrained by limited tissue availability and modest cohort sizes, and cryobiopsy sampling may underrepresent certain compartments, particularly immune populations. To address these limitations, we implemented a multi-layered validation strategy combining independent cohorts, complementary sampling approaches, and orthogonal technologies. The integration of single-nucleus and bulk transcriptomic data, together with spatial transcriptomics and immunohistochemical validation, enabled us to confirm the validity of our observations. The impressive consistency of our single cell findings across of samples obtained in two continents, two acquisition methods (cryobiopsy and VATs) and spanning 20 years, further enhances our confidence that our observations represent a true fundamental feature of early pulmonary fibrosis. Thus, the strength of our study lies in the integration of deeply phenotyped clinical cohorts spanning the earliest stages of disease. While prior studies have focused either on at-risk individuals or on patients with interstitial lung abnormalities, our design captures a broader and clinically relevant continuum, from asymptomatic familial IPF individuals to sporadic early disease with preserved lung function in multiple independent cohorts. This unique clinical framework enables the identification of convergent molecular and cellular features associated with disease initiation, rather than late-stage consequences. For decades, early pulmonary fibrosis has been somewhat of a clinical and mechanistic conundrum. Ten years ago, we learned that early interstitial findings should not be ignored, even minimal interstitial abnormalities frequently represent subclinical disease with a high risk of progression ^39^. Many studies have shown or confirmed that early pulmonary fibrosis is highly prevalent among first-degree relatives of patients with familial pulmonary fibrosis and is associated with functional decline and reduced survival ^27,29,36,40^. However, the nature of these early changes was not clear, where they similar to IPF, or did they represent an earlier inflammatory injury stage that is different form the full-blown disease? To these questions our results suggest an answer that needs substantial reckoning, cellularly and molecularly, early IPF is very similar to end stage IPF. While the extent of fibroblastic and macrophage changes is not as profound as in end stage disease, the epithelial changes are already present, suggesting that they are key features of disease. These findings establish a clinically grounded and biologically integrated framework for understanding disease initiation and highlight new opportunities for early diagnosis and therapeutic intervention aimed at preserving alveolar structure and function before irreversible fibrosis is established.

## Acknowledgments

We are grateful to all patients who participated in this study. Special thanks to Mei Zhong from the Yale Stem Cell Center genomics core facility who performed sequencing.

## Authors Contribution

## Material and Methods

### Selection of study cohort and study approval

Tissue samples of patients from Forli and Florence underwent cryobiopsy during ILD program at University of Florence and Forli, Italy, were used. Human lungs had been collected following local hospital ethical committee approval and informed patient consent. A secondary approval at Yale Institutional Review Board was obtained (IRB protocols: 2000030116, 2000030118). All histopathological diagnoses were independently reviewed by Prof. Homer (Yale School of Medicine). The associated transcriptomic datasets are currently being deposited in the Gene Expression Omnibus (GEO) repository.

Regarding the NIH cohort, approval was obtained from the National Heart, Lung, and Blood Institute and the National Human Genome Research Institute at the National Institutes of Health (Protocols 99-H-0068 and/or 04-HG-0211) and from the Baylor College of Medicine (IRB #H-46823). All subjects denied any history of lung disease at the time of study recruitment. Open-lung biopsies in these three subjects showed a pattern of UIP confirming a diagnosis of IPF. All the

### Tissue processing- Frozen tissue

Nuclei were extracted using the Nuclei Isolation kit (CG000505, 10X Genomics). Briefly and based on the manufacturer’s protocol and reagents, the tissue was dissociated on ice, centrifugated and washed. The pellet was resuspended and cellular debris were removed. Following another centrifugation step, nuclei were resuspended and counted.

### Single nuclei RNA sequencing

#### Single-cell barcoding, library preparation, and sequencing

Florence cohort - Around 20’000 nuclei (PCLS) were loaded on a Chip G with Chromium Single Cell 3′ v3.1 gel beads and reagents (3′ GEX v3.1, 10x Genomics). Final libraries were analyzed on an Agilent Bioanalyzer High Sensitivity DNA chip for qualitative control purposes. cDNA libraries were sequenced on a HiSeq 4000 Illumina platform aiming for 150 million reads per library and a sequencing configuration of 26 base pair (bp) on read1 and 98 bp on read2).

#### Processing Sequencing Data

Florence cohort - Basecalls were converted to reads with the software Cell Ranger’s (v4.0.0) implementation *mkfastq*. Multiple fastq files from the same library and read strand were catenated to individual read1 and read2 files before trimming. The program cutadapt (v3.0) was used to trim and filter reads prior to genome alignment. For all 10X Genomics 3-prime gene expression data, read2 files were subject to two passes of contaminant trimming (i) for the template switch oligo sequence (AAGCAGTGGTATCAACGCAGAGTACATGGG) anchored on the 5′ end and (ii) for poly(A) sequences anchored to the 3′ end. Trimmed reads were subject to genome alignment using the program STAR (v2.7.6a) and its STARsolo implementation. All libraries were mapped to the same STAR genome index of GRCh38.p13 using GENCODE’s human release 37 annotation scheme. Cell and UMI barcodes positions and manufacturer’s cell barcode whitelists were parameterized accordingly for each dataset.

#### snRNAseq Cleaning

Processed snRNAseq data cleaning, cell type labeling and exploratory analysis were performed in R (v4.3.3) using the package Seurat (v4.4.0). The top 2,000 variable genes were scaled across nuclei and used for principal component analysis (PCA); the top 50 PCs were used for a UMAP layout and louvain clustering.Clusters of low transcript abundance outliers which lacked defining features ,clusters with features representative of two or more otherwise-distinct populations ,clusters with a relatively high abundance of mitochondrial RNA, a high percent of spliced transcripts and/or extremely low abundances of the common nuclear-specific lncRNA MALAT1 were discarded. Clusters with quality nuclei profiles were assigned a cell type label.

#### snRNAseq Data Integration

To reduce unwanted signals from intersample variance, graph embeddings of the *in vivo* snRNAseq data were performed with data integration at the sample-level using Seurat’s implementation of reciprocal PCA (RPCA). Reference samples for integration were selected based on the diversity of cell types present and the level of detail/granularity observed during the cleaning process. Samples from the 4 IPF lungs (204_48_I,325_70_I,127_26_Ib,319_49_Ib) and 5 samples from 5 control lungs (248_111_D, 274_62_Db, 145_22_D, 304_71_Db, 222_68_D).

After splitting the data by either dataset or snRNAseq sample, the top 2,500 variable genes were selected from the reference samples using Seurat’s *SelectIntegrationFeatures* implementation. The number of transcripts and the percent of transcripts mitochondrial were both regressed out during the within-sample scaling and PCA steps of RPCA. Seurat’s *FindIntegrationAnchors* was run using the references d features, a *max.features* value of 250 and the *reduction* argument set to “rpca”. The final *IntegrateData* implementation was run under default parameters. Cell type marker genes were estimated using normalized empirical gene expression values, and calculations were performed for each dataset independently using the Seurat function *FindAllMarkers* with a minimum log2 fold change of 0.5.

#### NIH cohort

Single-nuclei data was mapped to the human genome build hg38 using Cell Ranger v8.0.0 (10x Genomics) Data was processed, checked for quality, and normalized using Scanpy^41^. Batch correction was performed at the sample level using SCVI ^42^. Cell types were determined using published gene markers based, ^30,32,43^. Final Uniform Manifold Approximation and Projection (UMAP) plots and cell counts were generated using Scanpy.

### CosmX-PPFE processing

5-µm sections were placed onto Superfrost Plus Micro Slides (VWR, 48311-703), dried, stored and then incubated overnight at 60 °C. Sections were dewaxed with xylene washes (or CitroSolv for normal tissues) and rehydrated through graded ethanol–water washes, followed by a final rinse in 1× PBS. Slides were dried at 60 °C for 5 minutes. Target Retrieval solution (CosMx FFPE slide preparation kit, RNA) was used in a pressure cooker at 100 °C for 15 minutes, followed by water, ethanol and air drying. Incubation frames were attached, and digestion buffer with 3 µg ml−1 Proteinase K (CosMx FFPE slide preparation kit, RNA) and 1× PBS was applied for 30 minutes. The slides were then rinsed, fiducials (0.001%) applied and fixed in 10% NBF washed with NBF stop buffer and 1× PBS. Next, 100 mM NHS acetate was applied for 15 minutes, followed by 2× saline-sodium citrate (SSC) washes. The CosMx Human Universal Cell Characterization RNA Panel targeting 950 human genes and a 50-target add-on panel set was denatured and cooled. A probe mix with RNase Inhibitor, Buffer R and nuclease-free water was applied for 17 hours at 37 °C. After incubation, slides were washed with 50% deionized formamide and 2× SSC and then in 2× SSC. DAPI stock (1:40) was applied for 15 minutes, washed with 1× PBS and stained for 1 hour with CD298/B2M, PanCK, CD45 and CD68 antibodies. Slides were washed in 1× PBS and stored in 2× SSC. The pre-bleaching followed Configuration C and cell segmentation Configuration A (except for the HCC TMA, Configuration E). Procedures and FOVs were consistent across all sites

#### Immunofluorescent staining

Formalin fixed, parrafin embedded tissue was cut at 5uM thickness and mounted on slides. Slides were deparaffinized and rehydrated: 5 minutes in xylene (repeated twice), 5 minutes in 100% ethanol, 5 minutes in 75% ethanol, 5 minutes in 50% ethanol and 5 minutes in PBS, Antigen retrieval was performed with a tris-based solution (Vector Labs; H-3301-250) at 95°C for 20 minutes and allowed to cool to room temperature. Slides were then treated with the autofluorescence quencher (Biotium; #23014) for 10 minutes per the manufacturer’s guidelines, then washed in PBS for 5 minutes, twice.

Tissue was exposed to a serum-free blocking agent (Dako; X0909) for one hour. Primary antibodies used for indirect labeling were diluted in the same block agent and incubated overnight at 4°C in a humid slide box. Slides were washed in PBS three times for 5 minutes each. Secondary antibodies were diluted 1:500 in a diluent of 2.5% donkey serum; slides were incubated with secondary antibodies for 1 hour at room temperature in a dark slide box; all subsequent steps shield the tissues from light to protect the fluorophores. Slides are then washed twice in PBS for 5 minutes each. Commercially preconjugated antibodies for direct labeling were diluted in 2.5% mouse serum and incubated for 30 minutes at room temperature to quench unbound secondary antibodies, then incubated on slides overnight in a humid slide box at 4°C. After washing slides twice in fresh PBS for 5 minutes each, another autofluorescence quenching treatment was performed (Vector Labs; SP-8500-15) for 5 minutes per the manufacturer’s instructions. Slides were subject to a final was step in PBS, three times for 5 minutes each. Coverslips were mounted with an aqueous mounting media containing Hoechst 33342 (Invitrogen; P36983) and left to cure overnight at room temperature. Slides were imaged with a Leica Stellaris 8 Falcon confocal microscope.

### Statistical analysis

Statistical analyses of cell frequencies were performed with the Kruskal-Wallis test. Post hoc intergroup analysis was conducted with the Wilcoxon rank sum test with Bonferroni correction for multiple testing on the cell type. Pearson correlation was used for assessing the relationship between the transcriptomic signatures of different cell types across different and independent cohorts. Statistical methods used throughout the snRNA-seq analysis pipeline are specified in the respective section of the methods. P values < 0.05 were considered statistically significant. Significance levels are indicated as follows: *P ≤ 0.05, **P ≤ 0.01, ***P ≤ 0.001, and ****P ≤ 0.0001. All statistical analyses were computed in R (v.4.2.1, The R Foundation).

