## Supplementary material for "Alveolar niche disruption and aberrant epithelial reprogramming are early hallmarks of idiopathic pulmonary fibrosis": supp

**Supplemental**

**Material and Methods**

Bulk RNA seq

Bulk RNA sequencing was performed on paraffin-embedded lung tissue samples. Raw gene-level count matrices were generated and imported into R for downstream analysis. Sample annotations, including disease condition, were provided in a separate metadata table. Differential gene expression analysis was conducted using the DESeq2 package (v1.51.5), (1). Analyses were performed separately for each comparison of interest, including IPF versus control. Genes with low expression were filtered prior to analysis, retaining genes with a total count of at least five reads across all samples in the comparison. Size factor estimation and normalization were performed using the DESeq2 median-of-ratios method to account for differences in library size and RNA composition. Dispersion estimates were obtained using empirical Bayes shrinkage, and model fitting was performed using the default DESeq2 pipeline. Differential expression results were extracted using Wald tests, and P values were adjusted for multiple testing using the Benjamini–Hochberg method to control the false discovery rate (FDR). Genes with an adjusted P value < 0.05 were considered significantly differentially expressed. The list of

For selected analyses and visualizations, an additional log2 fold-change threshold of ±1 (corresponding to a twofold change) was applied. Normalized counts were used for downstream visualization, including volcano plots and heatmaps. For heatmap visualization, normalized counts were log- using log1p transformation and scaled by gene (row-wise Z-score). Hierarchical clustering of genes and samples was performed using Euclidean distance and complete linkage.

BulkRNA seq Deconvolution

Deconvolution of the bulk RNA-seq data was conducted using CIBERSORT (2) with our Florence cohort sequencing (snRNA-seq) data as reference data, which contains 95,542 cells from 6 healthy control samples. Genes expressed in less than 3 cells were filtered out. Cells that captured expression of more than 200 genes and percentage of reads mapped to mitochondrial genome less than 30% were retained. Cell type annotation was retrieved from the original publication. The filtered annotated scRNA-seq data had 95,491 cells and 36,347 genes, which was uploaded onto CIBERSORTx server as reference scRNA-seq data, together with the bulk RNA-seq data for cell type deconvolution analysis. Signature gene matrix was built directly from the reference scRNA-seq data by CIBERSORTx under default settings.

*Spatial Transcriptomic analysis*

Neighborhood analysis

Giotto objects were generated for each field of view (FOV) from slides A, B, and C using the Giotto package (version 0.6.0) (3) in R. Each object included per-cell transcript coordinates derived from the transcript coordinate file and cell segmentation mask ‘tif’ file, as well as spatial centroid positions for each cell. Spatial network was then constructed for each FOV corresponding to IPF and control samples using the Delaunay triangulation method via createSpatialNetwork function in Giotto with parameters maximum_distance_delaunay = 200, a distance cutoff for defining nearest neighbors, and minimum_k = 0, setting the minimum number of nearest neighbors per cell. All other parameters were left at default values. These networks were converted into graph objects, and immediate neighbors for each cell were identified using ego function from the igraph package (version 1.4.2). An adjacency matrix was constructed from the neighbor relationships identified by ego function, and using known cell type annotations, cell-type-specific neighbor graphs were generated for both IPF and control conditions

Ligand receptor Analysis

Cell–cell communication analysis was performed using the CellChat R package (4) (v3) to infer ligand–receptor interactions from single-nucleus RNA sequencing (snRNA-seq) data. Analyses were conducted on a subset of cells corresponding to the alveolar niche, defined based on spatial transcriptomic annotation and including epithelial, endothelial, immune, and stromal populations.

For each condition, cells were first subset from the integrated Seurat object based on clinical annotation. To ensure robust inference and reduce bias from rare populations, cell types represented by fewer than 20 cells were excluded. To further balance cell-type representation, a downsampling strategy was applied, limiting each cell population to a maximum of 100 cells selected at random.

A CellChat object was then created using normalized gene expression data, with cell identities defined by annotated cell types. The CellChatDB.human database was used as the reference ligand–receptor interaction database. Data were preprocessed using the standard CellChat workflow, including identification of overexpressed genes and overexpressed ligand–receptor interactions.

Communication probabilities were computed using the computeCommunProb function based on raw expression values. Interactions supported by fewer than five cells were filtered out to reduce noise. Pathway-level communication probabilities were subsequently inferred using computeCommunProbPathway, and global communication networks were aggregated using aggregateNet.

To specifically interrogate communication within the alveolar niche, ligand–receptor inference was restricted to the subset of selected cell populations. The resulting interaction networks were visualized using circular plots (netVisual_circle) representing both the number and strength of inferred interactions. Pathway-specific communication was explored using hierarchical network visualization and ligand–receptor contribution analysis (netAnalysis_contribution).

Finally, inferred ligand–receptor interactions derived from snRNA-seq data were integrated with spatial transcriptomic data by mapping cell-type identities onto spatial coordinates, enabling the reconstruction of cell–cell communication networks within the tissue context.

**Supplemental Figure legend**

**Figure S1** Cohort characteristics and quality control metrics across datasets

**(A–C)** Florence cohort. **(A)** Distribution of clinical phenotypes within the Florence cohort, including early IPF (n = 14) and other interstitial lung diseases (ILDs; n = 8), with detailed subcategories (CTD-ILD, hypersensitivity pneumonitis, post-COVID fibrosis, pleuroparenchymal fibroelastosis, and smoking-related ILD). **(B)** Quality control metrics across samples, including UMI counts per cell, number of detected genes per cell, and percentage of mitochondrial reads, stratified by disease condition**. (C)** Proportion of major cell compartments (airway, alveolar, lymphoid, myeloid, stromal, and vascular endothelial) across conditions (Control, Early IPF, End-stage IPF, and other ILDs), highlighting disease-associated shifts in cellular composition.

**(D–E) NIH cohort (D)** Relative abundance of major lung cell compartments across Control, early familial pulmonary fibrosis (early FPF), and IPF samples. **(E)** Quality control metrics across individual samples, including UMI counts, number of detected genes, and mitochondrial read percentages, demonstrating consistent data quality across phenotypes.

**(F–G) Forlì cohort.** **(F)** Distribution of clinical phenotypes in the Forlì cohort (n = 24), including early IPF (n = 8), controls (n = 4), and other ILDs (n = 12), with detailed etiologies. **(G)** Quality control metrics across samples, including UMI counts per cell and gene detection levels, stratified by condition, confirming the robustness and comparability of sequencing data across cohorts.

**Figure S2 Supplementary Cellular landscape in the NIH validation cohort**

**(A)** UMAP representation of single-nucleus RNA-seq data. Dimensionality reduction of the NIH cohort highlighting major lung cell populations, including epithelial, immune, stromal, and vascular compartments. Cell types are annotated and color-coded based on canonical marker gene expression**.(B)** UMAP colored by condition. Projection of cells colored by phenotype (Control, early familial pulmonary fibrosis (early FPF), and idiopathic pulmonary fibrosis (IPF)), illustrating the distribution of disease states across cellular populations**.(C)** Heatmap of cell-type–specific gene expression. Normalized expression of selected marker genes across all identified cell types, displayed as scaled values. This analysis confirms robust annotation of major lung cell populations and highlights distinct transcriptional programs across epithelial, immune, and stromal compartments**.(D)** Correlation matrix of transcriptional profiles. Pairwise correlation analysis of gene expression signatures across cell types, revealing strong transcriptional coherence within lineages and expected relationships between epithelial, immune, and stromal compartments.

**Figure S3 Transcriptional programs and inferred cellular composition in the Forlì cohort**

**(A)** Heatmap of scaled gene expression (Z-scores).Hierarchical clustering of differentially expressed genes across samples, displayed as Z-score–scaled expression values. This representation highlights relative up- and downregulation patterns between control and IPF samples, revealing distinct transcriptional programs associated with fibrotic remodeling. Annotation bars indicate sample identity and condition. **(B)** Volcano plot of differential gene expression. Differential expression analysis comparing IPF and control samples. Each point represents a gene plotted by log₂ fold change and –log₁₀ adjusted p-value. Significantly upregulated genes in IPF are highlighted in red, whereas downregulated genes are shown in blue. Selected genes of interest are annotated, illustrating key pathways associated with fibrotic remodeling**.(C)** Inferred cell-type composition by deconvolution. 
Estimated proportions of major lung cell populations derived from bulk transcriptomic data using a deconvolution approach. Boxplots represent the distribution of inferred cell-type proportions across samples, stratified by condition (Control in blue, IPF in red). These results reveal disease-associated shifts in cellular composition, including alterations in epithelial and stromal compartments.

**Figure S4** **Spatial architecture and intercellular communication within the fibrotic alveolar niche**

**(A)** Representative airway regions**.**Multiplex imaging of airway regions in the Forlì cohort, showing epithelial and stromal organization. Marker staining (including epithelial and immune markers) highlights airway architecture. Insets indicate regions of interest used for downstream analyses, with corresponding cell segmentation and annotation. **(B)** UMAP representation of spatial transcriptomic data. Dimensionality reduction of spatially resolved cells identifying major lung cell populations, including epithelial, immune, and stromal compartments. Cell types are annotated based on marker expression. **(C)** UMAP colored by condition. Projection of cells colored by condition (Control vs IPF), illustrating disease-associated shifts in cellular distribution**.(D)** Delaunay-based spatial neighborhood network**.** Delaunay triangulation was used to construct spatial neighborhood networks from segmented cells, enabling the identification of physically adjacent cell populations within control and IPF tissue regions. Nodes represent cell types and edges reflect spatial proximity-based interactions, highlighting remodeling of local cellular neighborhoods in IPF**.(E)** Cell–cell neighborhood analysis illustrating spatial organization of the alveolar niche in control and IPF lungs. Network diagrams show the relative abundance of neighboring cell types surrounding AT1 epithelial cells, highlighting increased interactions with fibroblasts, macrophages and aberrant epithelial states in IPF. (**F)** Global network of predicted cell–cell interactions between major cell populations identified within the alveolar niche based on spatial transcriptomic data, including alveolar epithelial cells, immune cells, endothelial cells, and fibroblasts. For this analysis, cells composing the alveolar niche were subset and ligand–receptor interactions were inferred using single-nucleus RNA sequencing data, and subsequently projected onto the spatial dataset. Edge thickness represents the number of inferred interactions, highlighting the complexity of intercellular communication in IPF**.(G)** Predicted ligand–receptor signaling pathways mediating communication between cell types within the IPF alveolar niche, including collagen, PDGF, WNT, CDH, and TGFβ signaling pathways. Pathway inference was performed on the subset of cells defining the alveolar niche using single-nucleus RNA sequencing data. Heat maps indicate the relative contribution of sender, receiver, and mediator cell populations for each pathway.

**Tables**

Table Panel probe gene list – COSMx spatial transcriptomic analysis

| ABL1 | ABL2 | ACE | ACE2 | ACKR1 | ACKR3 | ACKR4 | ACTA2 | ACTG2 | ACVR1 |
| --- | --- | --- | --- | --- | --- | --- | --- | --- | --- |
| ACVR1B | ACVR2A | ACVRL1 | ADGRA2 | ADGRA3 | ADGRB2 | ADGRB3 | ADGRD1 | ADGRE1 | ADGRE2 |
| ADGRE5 | ADGRF1 | ADGRF3 | ADGRF4 | ADGRF5 | ADGRG1 | ADGRG2 | ADGRG3 | ADGRG5 | ADGRG6 |
| ADGRL1 | ADGRL2 | ADGRL4 | ADGRV1 | ADIPOQ | ADIRF | ADM2 | ADORA2A | AGR2 | AHI1 |
| AHR | AKT1 | ALCAM | ANGPT1 | ANGPT2 | ANGPT4 | ANGPTL1 | ANXA1 | ANXA2 | ANXA4 |
| APOA1 | APOB | APOD | APP | AQP3 | AR | AREG | ARF1 | ARG1 | ARHGDIB |
| ARTN | ATF3 | ATG10 | ATG12 | ATG5 | ATM | ATR | AXL | AZGP1 | AZU1 |
| B2M | B3GNT7 | BAG3 | BATF3 | BAX | BCL2 | BCL2L1 | BECN1 | BEST1 | BGN |
| BID | BIRC5 | BMP1 | BMP2 | BMP3 | BMP4 | BMP5 | BMP6 | BMP7 | BMPR1A |
| BMPR2 | BMX | BRCA1 | BST1 | BST2 | BTG1 | BTK | C11orf96 | C1QA | C1QB |
| C1QC | C5AR2 | C9orf16 | CALB1 | CALD1 | CALM1 | CALM2 | CALM3 | CAMP | CASP3 |
| CASP8 | CASR | CAV1 | CCL11 | CCL13 | CCL15 | CCL18 | CCL19 | CCL2 | CCL20 |
| CCL21 | CCL23 | CCL26 | CCL28 | CCL3 | CCL3L3 | CCL4 | CCL4L2 | CCL5 | CCL7 |
| CCL8 | CCND1 | CCR1 | CCR10 | CCR2 | CCR5 | CCR7 | CCRL2 | CD14 | CD163 |
| CD164 | CD19 | CD2 | CD209 | CD24 | CD27 | CD274 | CD276 | CD28 | CD300A |
| CD33 | CD34 | CD36 | CD37 | CD38 | CD3D | CD3E | CD3G | CD4 | CD40 |
| CD40LG | CD44 | CD47 | CD48 | CD52 | CD53 | CD55 | CD58 | CD59 | CD5L |
| CD63 | CD68 | CD69 | CD70 | CD74 | CD79A | CD80 | CD81 | CD83 | CD84 |
| CD86 | CD8A | CD8B | CD9 | CDH1 | CDH11 | CDH5 | CDKN1A | CDKN3 | CEACAM1 |
| CEACAM6 | CELSR1 | CELSR2 | CENPF | CFD | CFLAR | CHEK1 | CHEK2 | CHGA | CHI3L1 |
| CIDEA | CIITA | CLCF1 | CLDN4 | CLEC10A | CLEC12A | CLEC14A | CLEC1A | CLEC2B | CLEC2D |
| CLEC4A | CLEC4D | CLEC4E | CLEC5A | CLEC7A | CLOCK | CLU | CMKLR1 | CNTFR | COL11A1 |
| COL12A1 | COL14A1 | COL15A1 | COL16A1 | COL17A1 | COL18A1 | COL1A1 | COL1A2 | COL21A1 | COL27A1 |
| COL3A1 | COL4A1 | COL4A2 | COL4A5 | COL5A1 | COL5A2 | COL5A3 | COL6A1 | COL6A2 | COL6A3 |
| COL8A1 | COL9A1 | COL9A2 | COL9A3 | COTL1 | CPA3 | CPB1 | CRIP1 | CRP | CRYAB |
| CSF1 | CSF1R | CSF2 | CSF2RA | CSF2RB | CSF3 | CSF3R | CSHL1 | CSK | CST7 |
| CTLA4 | CTNNB1 | CTSG | CTSW | CUZD1 | CX3CL1 | CX3CR1 | CXCL1 | CXCL10 | CXCL12 |
| CXCL14 | CXCL16 | CXCL17 | CXCL2 | CXCL3 | CXCL5 | CXCL6 | CXCL8 | CXCL9 | CXCR1 |
| CXCR2 | CXCR3 | CXCR4 | CXCR5 | CXCR6 | CYP19A1 | CYP1B1 | CYSTM1 | CYTOR | DCN |
| DDC | DDIT3 | DDR1 | DDR2 | DDX58 | DHRS2 | DLL1 | DMBT1 | DNMT1 | DNMT3A |
| DNTT | DPP4 | DST | DUSP1 | DUSP2 | DUSP4 | DUSP5 | DUSP6 | EFNA1 | EFNA4 |
| EFNA5 | EFNB1 | EFNB2 | EFNB3 | EGF | EGFR | EIF5A | ELANE | EMP3 | ENG |
| ENTPD1 | EOMES | EPCAM | EPHA2 | EPHA3 | EPHA4 | EPHA7 | EPHB2 | EPHB3 | EPHB4 |
| EPHB6 | EPOR | ERBB2 | ERBB3 | ESAM | ESR1 | ETS1 | ETV4 | ETV5 | EZH2 |
| EZR | FABP4 | FABP5 | FAS | FASLG | FASN | FCER1G | FCGBP | FCGR3A | FCRLA |
| FES | FFAR2 | FFAR3 | FFAR4 | FGF1 | FGF12 | FGF13 | FGF18 | FGF2 | FGF7 |
| FGF9 | FGFR1 | FGFR2 | FGFR3 | FGG | FGR | FKBP11 | FLT1 | FLT3LG | FN1 |
| FOS | FOXF1 | FOXP3 | FPR1 | FYB1 | FYN | FZD1 | FZD3 | FZD4 | FZD5 |
| FZD6 | FZD7 | FZD8 | G6PC2 | G6PD | GADD45B | GAS6 | GATA3 | GC | GCG |
| GDF10 | GDF15 | GDF3 | GDF6 | GDF9 | GDNF | GLUD1 | GLUL | GNLY | GPBAR1 |
| GPER1 | GPNMB | GPR183 | GPX1 | GPX3 | GSN | GSTP1 | GZMA | GZMB | GZMH |
| GZMK | H2AZ1 | H4C3 | HAVCR2 | HBA1 | HBB | HCAR2 | HCAR3 | HCK | HCST |
| HDAC1 | HDAC11 | HDAC3 | HDAC4 | HDAC5 | HGF | HIF1A | HILPDA | HLA-A | HLA-B |
| HLA-C | HLA-DPA1 | HLA-DPB1 | HLA-DQA1 | HLA-DQB1 | HLA-DRA | HLA-DRB1 | HLA-DRB5 | HLA-E | HMGB2 |
| HMGN2 | HPGDS | HSD17B2 | HSD3B2 | HSP90AA1 | HSP90AB1 | HSP90B1 | HSPA1A | HSPA1B | HSPB1 |
| HTT | IAPP | ICAM1 | ICAM2 | ICAM3 | ICOS | ICOSLG | IDO1 | IER3 | IFI27 |
| IFIH1 | IFIT1 | IFITM1 | IFITM3 | IFNA1 | IFNAR1 | IFNAR2 | IFNB1 | IFNG | IFNGR1 |
| IFNGR2 | IFNL2 | IFNL3 | IGF1 | IGF1R | IGF2 | IGF2R | IGFBP3 | IGFBP5 | IGFBP6 |
| IGFBP7 | IGHA1 | IGHD | IGHG1 | IGHG2 | IGHM | IGKC | IL10 | IL10RA | IL10RB |
| IL11 | IL11RA | IL12A | IL12B | IL12RB1 | IL12RB2 | IL13RA1 | IL15 | IL15RA | IL16 |
| IL17A | IL17B | IL17D | IL17RA | IL17RB | IL17RE | IL18 | IL18R1 | IL1A | IL1B |
| IL1R1 | IL1R2 | IL1RAP | IL1RL1 | IL1RN | IL2 | IL20 | IL20RA | IL22RA1 | IL23A |
| IL24 | IL27RA | IL2RA | IL2RB | IL2RG | IL32 | IL33 | IL34 | IL36G | IL3RA |
| IL4R | IL6 | IL6R | IL6ST | IL7 | IL7R | INHA | INHBA | INHBB | INS |
| INSR | IRF3 | IRF4 | ITGA1 | ITGA2 | ITGA3 | ITGA5 | ITGA6 | ITGA9 | ITGAE |
| ITGAL | ITGAM | ITGAV | ITGAX | ITGB1 | ITGB2 | ITGB4 | ITGB5 | ITGB6 | ITGB8 |
| ITK | ITM2A | JAG1 | JAK1 | JAK2 | JCHAIN | JUN | JUNB | KDR | KIT |
| KITLG | KLF2 | KLK3 | KLRB1 | KLRK1 | KRAS | KRT1 | KRT10 | KRT13 | KRT14 |
| KRT15 | KRT16 | KRT17 | KRT18 | KRT19 | KRT20 | KRT23 | KRT24 | KRT4 | KRT5 |
| KRT6A | KRT6B | KRT6C | KRT7 | KRT8 | KRT80 | KRT86 | LAG3 | LAIR1 | LAMP2 |
| LAMP3 | LCN2 | LDLR | LEFTY1 | LEFTY2 | LEP | LGALS1 | LGALS3 | LGALS3BP | LGALS9 |
| LIF | LIFR | LINC02446 | LMNA | LPAR5 | LTB | LTBR | LTF | LUM | LY6D |
| LY75 | LYN | LYZ | MAF | MALAT1 | MAML2 | MAP1LC3B | MAPK13 | MAPK14 | MARCO |
| MECOM | MEG3 | MERTK | MET | MGP | MIF | MKI67 | MMP1 | MMP10 | MMP12 |
| MMP14 | MMP16 | MMP19 | MMP2 | MMP3 | MMP7 | MMP8 | MMP9 | MPO | MRC1 |
| MRC2 | MS4A1 | MS4A4A | MSMB | MST1R | MT1X | MT2A | MTOR | MTRNR2L1 | MX1 |
| MXRA8 | MYC | MYH11 | MYL9 | MZB1 | MZT2A | NANOG | NCR1 | NDRG1 | NEAT1 |
| NFKB1 | NFKBIA | NGFR | NKG7 | NLRC4 | NLRC5 | NLRP1 | NLRP2 | NLRP3 | NOD2 |
| NOSIP | NOTCH1 | NOTCH2 | NOTCH3 | NPPB | NPPC | NPR1 | NPR2 | NPR3 | NR1H2 |
| NR1H3 | NR1H4 | NR3C1 | NRG1 | NRG4 | NRIP3 | NRXN1 | NRXN3 | NTRK2 | OAS1 |
| OAS2 | OAS3 | OASL | OLFM4 | OLR1 | OSM | OSMR | OXER1 | OXGR1 | P2RX5 |
| P2RY12 | PARP1 | PCNA | PDCD1 | PDCD1LG2 | PDGFA | PDGFB | PDGFC | PDGFD | PDGFRA |
| PDGFRB | PECAM1 | PF4 | PGF | PGR | PHLDA2 | PIGR | PLA2R1 | PLAC8 | PNOC |
| POU5F1 | PPARA | PPARD | PPARG | PPBP | PRF1 | PROK2 | PROKR1 | PRSS2 | PSAP |
| PSCA | PTGDR2 | PTGDS | PTGES | PTGES2 | PTGES3 | PTGIS | PTGS1 | PTGS2 | PTHLH |
| PTK2 | PTK6 | PTPRC | PTPRCAP | PTTG1 | QRFPR | RAC1 | RAC2 | RAD51 | RAMP1 |
| RAMP2 | RAMP3 | RARA | RARB | RARG | RARRES1 | RARRES2 | RB1 | RBPJ | REG1A |
| RELA | RELT | RGCC | RGS1 | RGS2 | RGS5 | RNF43 | ROR1 | RORA | RPL21 |
| RPL22 | RPL32 | RPL34 | RPL37 | RPS4Y1 | RSPO1 | RSPO2 | RSPO3 | RUNX3 | RXRA |
| RXRB | RYK | S100A10 | S100A2 | S100A4 | S100A6 | S100A8 | S100A9 | S100B | S100P |
| SAA1 | SAA2 | SAT1 | SCG5 | SCGB3A1 | SEC23A | SEC61G | SELENOP | SELL | SELPLG |
| SERPINA1 | SERPINA3 | SERPINB5 | SERPINH1 | SFN | SIGIRR | SLC2A1 | SLC2A4 | SLC40A1 | SLPI |
| SMAD2 | SMAD3 | SMAD4 | SMARCB1 | SMO | SNAI1 | SNAI2 | SOD1 | SOD2 | SOSTDC1 |
| SOX2 | SOX4 | SOX9 | SPARCL1 | SPINK1 | SPOCK2 | SPP1 | SPRY2 | SPRY4 | SQSTM1 |
| SRC | SREBF1 | SRGN | SST | ST6GAL1 | ST6GALNAC3 | STAT1 | STAT3 | STAT4 | STAT5A |
| STAT5B | STAT6 | STMN1 | SUCNR1 | SYK | TACSTD2 | TAGLN | TAP1 | TAP2 | TBX21 |
| TCL1A | TEK | TFEB | TGFB1 | TGFB2 | TGFB3 | TGFBR1 | TGFBR2 | THBS1 | THBS2 |
| TIE1 | TIGIT | TIMP1 | TLR1 | TLR2 | TLR3 | TLR4 | TLR5 | TLR7 | TLR8 |
| TM4SF1 | TNF | TNFAIP6 | TNFRSF10A | TNFRSF10B | TNFRSF10D | TNFRSF11A | TNFRSF11B | TNFRSF12A | TNFRSF13B |
| TNFRSF14 | TNFRSF17 | TNFRSF18 | TNFRSF19 | TNFRSF1A | TNFRSF1B | TNFRSF21 | TNFRSF4 | TNFRSF9 | TNFSF10 |
| TNFSF12 | TNFSF13B | TNFSF14 | TNFSF15 | TNFSF18 | TNFSF4 | TNFSF8 | TNFSF9 | TOP2A | TOX |
| TP53 | TPM1 | TPM2 | TPSAB1 | TPSB2 | TSC22D1 | TSHZ2 | TSLP | TTR | TUBB |
| TUBB4B | TWIST1 | TWIST2 | TXK | TYK2 | TYMS | TYROBP | UBE2C | UCP1 | UPK3A |
| VCAM1 | VCAN | VEGFA | VEGFB | VEGFC | VEGFD | VHL | VIM | VPREB3 | VSIR |
| VTN | VWF | WIF1 | WNT10B | WNT11 | WNT2 | WNT2B | WNT3 | WNT7A | WNT7B |
| WNT9A | XBP1 | XCL1 | XCL2 | YBX3 | YES1 | ZFP36 | NegPrb3 | NegPrb5 | NegPrb6 |
| NegPrb7 | NegPrb8 | NegPrb9 | NegPrb10 | NegPrb11 | NegPrb12 | NegPrb13 | NegPrb14 | NegPrb15 | NegPrb16 |
| NegPrb18 | NegPrb19 | NegPrb20 | NA | NA | NA | NA | NA | NA | NA |

Table S2 Differential gene expression analysis between Early IPF and controls using DESeq2

| Genes | baseMean | log2FoldChange | lfcSE | stat | pvalue | padj |
| --- | --- | --- | --- | --- | --- | --- |
| LTA | 66.99763166 | 8.613006752 | 1.65500661 | 5.20421289 | 1.95E-07 | 0.00058953 |
| MMP7 | 64.49465686 | 8.560252814 | 2.08634408 | 4.10299187 | 4.08E-05 | 0.02372491 |
| TNFRSF13B | 48.20836384 | 8.139116437 | 1.92572297 | 4.22652508 | 2.37E-05 | 0.01840774 |
| DEPDC1B | 42.96476086 | 7.973372716 | 1.97191152 | 4.04347387 | 5.27E-05 | 0.02450899 |
| KRT24 | 68.63898553 | 7.967834527 | 2.04959642 | 3.88751387 | 0.00010128 | 0.03150724 |
| KRT16P2 | 255.7589257 | 7.830577413 | 1.2706625 | 6.16259424 | 7.16E-10 | 1.33E-05 |
| CD1B | 35.81191961 | 7.711146254 | 2.02866933 | 3.80108584 | 0.00014406 | 0.03806578 |
| IL1RN | 260.4788712 | 7.536153741 | 1.45480919 | 5.18016643 | 2.22E-07 | 0.00058953 |
| TMEM95 | 24.0876358 | 7.139077434 | 1.74609074 | 4.0886062 | 4.34E-05 | 0.02376001 |
| EREG | 23.06622133 | 7.077781346 | 1.81791887 | 3.8933428 | 9.89E-05 | 0.03150724 |
| C20orf195 | 60.44853341 | 6.799235267 | 1.62278209 | 4.1898634 | 2.79E-05 | 0.01985416 |
| LOC440461 | 88.4686526 | 6.715863171 | 1.21645215 | 5.52086094 | 3.37E-08 | 0.00020932 |
| CCL13 | 56.88525408 | 6.700403846 | 1.65380514 | 4.05150745 | 5.09E-05 | 0.02428957 |
| CLDN16 | 54.37847266 | 6.631819022 | 1.7640182 | 3.75949581 | 0.00017026 | 0.04225757 |
| C15orf48 | 58.55396522 | 6.609440645 | 1.62809318 | 4.05962063 | 4.92E-05 | 0.02428957 |
| ASF1B | 26.58744944 | 6.504945523 | 1.70098256 | 3.82422823 | 0.00013118 | 0.03674182 |
| APOBEC3B-AS1 | 44.07030593 | 6.454735738 | 1.45321878 | 4.44168202 | 8.93E-06 | 0.00977379 |
| EPHX3 | 42.31512258 | 6.436601601 | 1.63751005 | 3.93072494 | 8.47E-05 | 0.0294898 |
| TRIM9 | 60.19298059 | 5.972255342 | 1.50507454 | 3.96807945 | 7.25E-05 | 0.02869648 |
| BHLHA15 | 93.44688935 | 5.663347743 | 1.30184212 | 4.35025698 | 1.36E-05 | 0.0140624 |
| AGRP | 89.82902889 | 5.629551283 | 1.35002118 | 4.16997258 | 3.05E-05 | 0.01985416 |
| MYBL2 | 134.7125375 | 5.474257115 | 1.28289106 | 4.26712546 | 1.98E-05 | 0.01755198 |
| BEX5 | 25.35555173 | 5.465518304 | 1.28828295 | 4.24248285 | 2.21E-05 | 0.01789148 |
| TMEM213 | 158.208196 | 5.090950769 | 0.97490963 | 5.22197198 | 1.77E-07 | 0.00058953 |
| ADGRE1 | 108.3852763 | 4.989758093 | 1.21828007 | 4.09573974 | 4.21E-05 | 0.02373821 |
| MNX1 | 35.87476408 | 4.807976861 | 1.22530897 | 3.9238894 | 8.71E-05 | 0.0294898 |
| KRT16P1 | 75.10040888 | 4.685193718 | 1.12658246 | 4.15876677 | 3.20E-05 | 0.01985416 |
| DIO2 | 89.16071889 | 4.59192901 | 1.10193712 | 4.1671425 | 3.08E-05 | 0.01985416 |
| SV2A | 114.8673617 | 4.199408297 | 0.80253683 | 5.23266742 | 1.67E-07 | 0.00058953 |
| RSPO2 | 53.02451942 | 4.175179732 | 1.0944653 | 3.81481234 | 0.00013629 | 0.03730842 |
| HIF1A-AS1 | 253.9453424 | 3.783621087 | 0.8337551 | 4.53804851 | 5.68E-06 | 0.00660567 |
| C2CD4B | 114.0920736 | 3.701981841 | 0.9745405 | 3.7986947 | 0.00014546 | 0.03806578 |
| HIST1H2AK | 124.8216784 | 3.550831631 | 0.74853361 | 4.74371705 | 2.10E-06 | 0.00300463 |
| HPSE2 | 186.9918968 | 3.509450173 | 0.62038166 | 5.65692119 | 1.54E-08 | 0.00014344 |
| PTGS2 | 298.4352177 | 3.473260393 | 0.91505346 | 3.79569124 | 0.00014723 | 0.03806578 |
| TMC3 | 128.3200887 | 3.263395181 | 0.82690706 | 3.94650784 | 7.93E-05 | 0.0294898 |
| IGLL5 | 8783.549366 | 3.186576038 | 0.85817734 | 3.71319061 | 0.00020466 | 0.0470345 |
| CLDN3 | 194.2041078 | 3.110416215 | 0.64656835 | 4.8106534 | 1.50E-06 | 0.0028004 |
| CPXM1 | 416.0115655 | 2.91609688 | 0.72912602 | 3.99944155 | 6.35E-05 | 0.02748619 |
| CDKN1A | 749.1883819 | 2.891535726 | 0.72777829 | 3.97309974 | 7.09E-05 | 0.02869648 |
| FER1L4 | 795.1282159 | 2.427366967 | 0.65484661 | 3.70677183 | 0.00020992 | 0.04765393 |
| HSH2D | 434.5404642 | 2.390288745 | 0.63645916 | 3.75560427 | 0.00017292 | 0.04235494 |
| MIR374B | 113.4517908 | 2.344841537 | 0.58355282 | 4.01821644 | 5.86E-05 | 0.02599023 |
| MIR374C | 113.4517908 | 2.344841537 | 0.58355282 | 4.01821644 | 5.86E-05 | 0.02599023 |
| SPHK1 | 662.9196754 | 2.221514045 | 0.47231294 | 4.70347912 | 2.56E-06 | 0.00340077 |
| VWA1 | 546.7118676 | 2.180026574 | 0.57611872 | 3.78398846 | 0.00015434 | 0.03935542 |
| HIST1H4K | 795.4970561 | 2.169345187 | 0.54388619 | 3.98860131 | 6.65E-05 | 0.02749395 |
| HIST1H4J | 797.5387091 | 2.141919608 | 0.54497958 | 3.93027498 | 8.48E-05 | 0.0294898 |
| CRISPLD2 | 1364.599787 | 2.074042221 | 0.48359409 | 4.288808 | 1.80E-05 | 0.01671949 |
| STAP2 | 364.404407 | 2.028384248 | 0.4241637 | 4.78207885 | 1.73E-06 | 0.00293595 |
| FBLIM1 | 764.6866207 | 1.934913164 | 0.47613198 | 4.06381683 | 4.83E-05 | 0.02428957 |
| TNC | 5658.844892 | 1.870599696 | 0.3926709 | 4.76378488 | 1.90E-06 | 0.0029473 |
| TTC39C | 428.9101752 | 1.797047894 | 0.41540319 | 4.32603295 | 1.52E-05 | 0.01487424 |
| FUCA2 | 564.8524286 | 1.643819941 | 0.4186105 | 3.92684832 | 8.61E-05 | 0.0294898 |
| LOC388242 | 818.7782939 | 1.581400065 | 0.41373607 | 3.82224366 | 0.00013224 | 0.03674182 |
| LOC613038 | 818.7782939 | 1.581400065 | 0.41373607 | 3.82224366 | 0.00013224 | 0.03674182 |
| CAMK1D | 1133.75006 | 1.494612666 | 0.38397381 | 3.89248601 | 9.92E-05 | 0.03150724 |
| CHPF | 1056.268146 | 1.481958421 | 0.37890544 | 3.91115637 | 9.19E-05 | 0.03053368 |
| EMILIN1 | 2091.772814 | 1.266635021 | 0.31240261 | 4.05449563 | 5.02E-05 | 0.02428957 |
| PFKP | 2229.003761 | 1.203397124 | 0.256821 | 4.68574267 | 2.79E-06 | 0.00346173 |
| SLC39A4 | 499.9454649 | 1.201954225 | 0.29171676 | 4.12027825 | 3.78E-05 | 0.02272322 |
| LRRC8C | 684.2141949 | 1.19203199 | 0.30748859 | 3.87667065 | 0.0001059 | 0.0323155 |
| TRAM2 | 1048.337842 | 1.134300381 | 0.29817929 | 3.80408846 | 0.00014233 | 0.03806578 |
| VSIG10 | 1147.45064 | -1.112583429 | 0.28345967 | -3.9250149 | 8.67E-05 | 0.0294898 |
| DNAJC19 | 776.7840664 | -1.640749147 | 0.43779773 | -3.7477333 | 0.00017844 | 0.04244014 |
| KLF15 | 625.8264174 | -1.783765431 | 0.45210892 | -3.945433 | 7.97E-05 | 0.0294898 |
| TMEM133 | 105.2953109 | -1.814919829 | 0.46893599 | -3.8702933 | 0.0001087 | 0.03263765 |
| TBC1D4 | 6265.821071 | -1.933352919 | 0.47416245 | -4.0774062 | 4.55E-05 | 0.02422125 |
| SH3RF2 | 2268.671717 | -1.944172128 | 0.52208668 | -3.7238493 | 0.00019621 | 0.04565517 |
| NEDD4 | 2436.66798 | -2.041418019 | 0.54504768 | -3.7453935 | 0.00018011 | 0.04244014 |
| FILIP1 | 2250.461315 | -2.333327384 | 0.55880568 | -4.1755613 | 2.97E-05 | 0.01985416 |
| CLCN4 | 769.2935102 | -2.500092937 | 0.62646561 | -3.9907904 | 6.59E-05 | 0.02749395 |
| WDR62 | 1087.726248 | -2.529324208 | 0.64027438 | -3.9503755 | 7.80E-05 | 0.0294898 |
| GHR | 1194.541753 | -2.606589745 | 0.61299368 | -4.2522294 | 2.12E-05 | 0.01789148 |
| UBE2D1 | 549.3661757 | -2.645261617 | 0.68818946 | -3.8437985 | 0.00012114 | 0.03523601 |
| POTEF | 1513.01695 | -3.344506842 | 0.68191798 | -4.9045588 | 9.36E-07 | 0.00193674 |
| KLHL33 | 294.0675743 | -3.362933737 | 0.89171422 | -3.7713134 | 0.00016239 | 0.04085001 |
| GBAS | 1995.147834 | -3.472673751 | 0.83418557 | -4.1629511 | 3.14E-05 | 0.01985416 |
| LRRC66 | 176.1161346 | -3.914813106 | 1.01399681 | -3.8607746 | 0.00011303 | 0.03339713 |
| KLHL30 | 591.1096626 | -4.203013373 | 1.0813424 | -3.8868479 | 0.00010155 | 0.03150724 |
| PRKAG3 | 983.0923752 | -4.734656114 | 1.26297927 | -3.7487995 | 0.00017768 | 0.04244014 |
| LRRC38 | 112.5021972 | -10.59784151 | 2.11471925 | -5.011465 | 5.40E-07 | 0.00125691 |
