## Supplementary figures and images for "Alveolar niche disruption and aberrant epithelial reprogramming are early hallmarks of idiopathic pulmonary fibrosis"

### sup1

#### Florence cohort

A

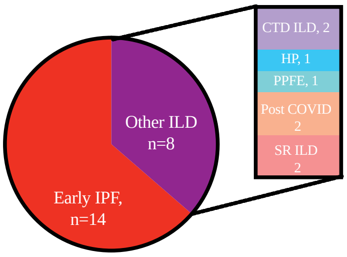


B

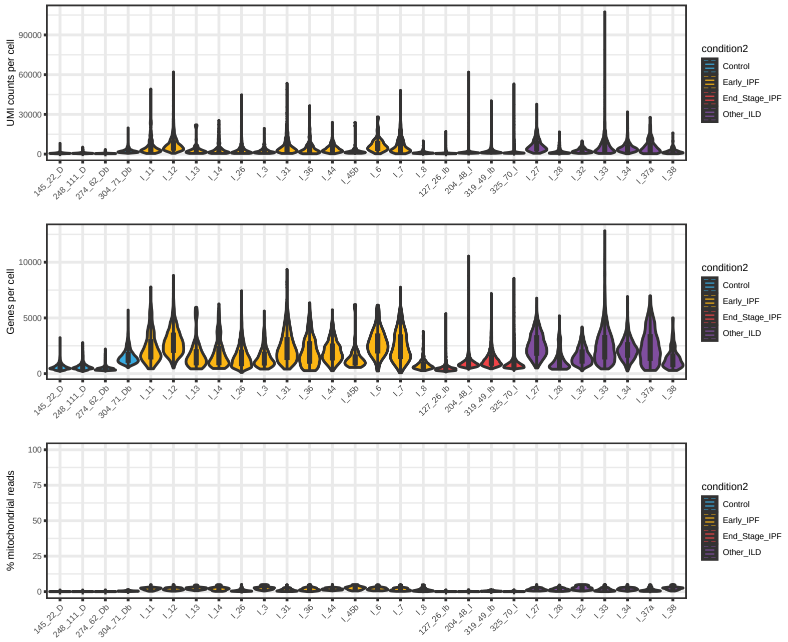


C

Total=22

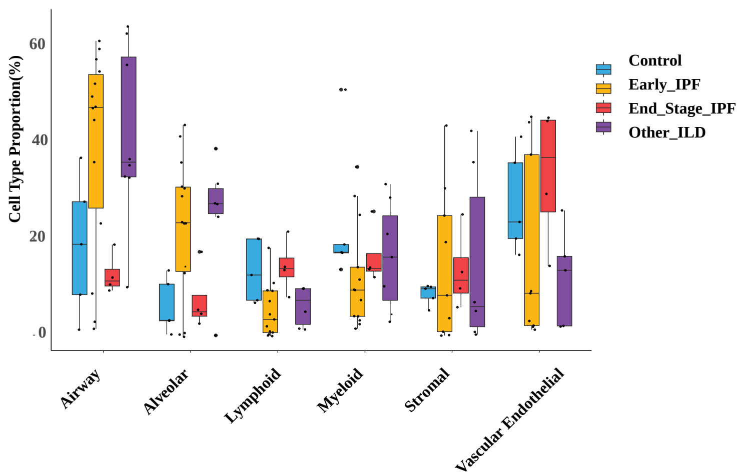


#### NIH cohort, n=9

D

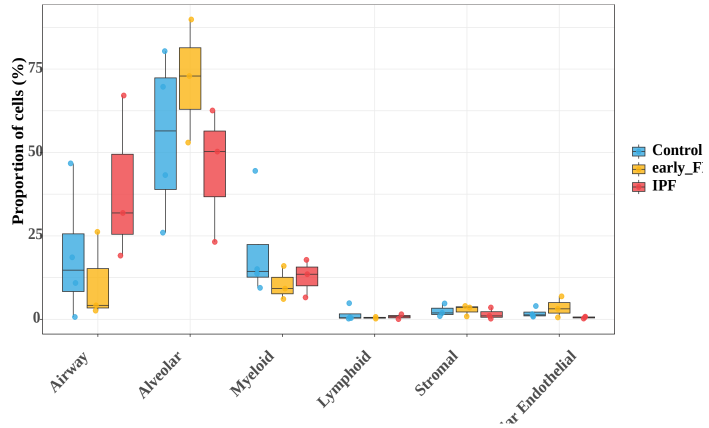


E

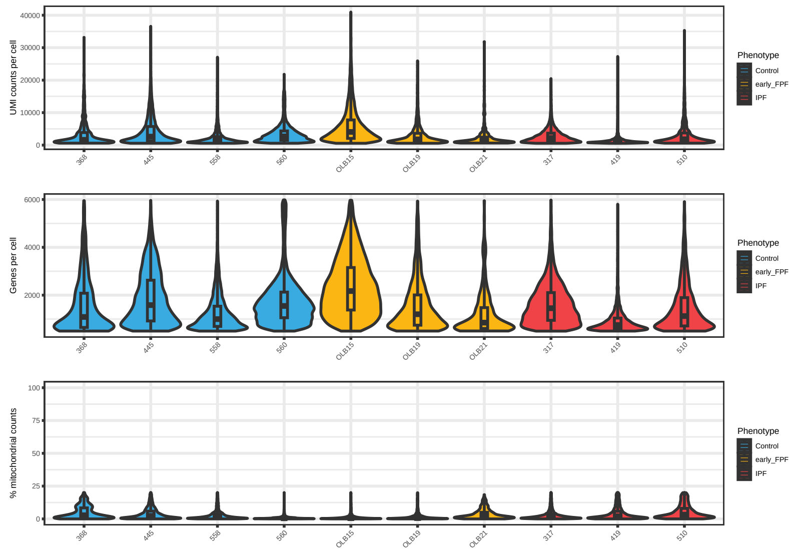


#### Forli cohort

F

Total=24

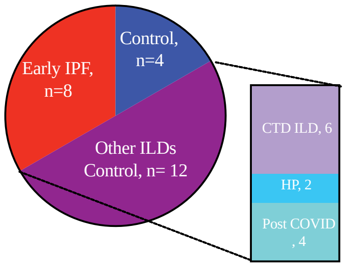


G

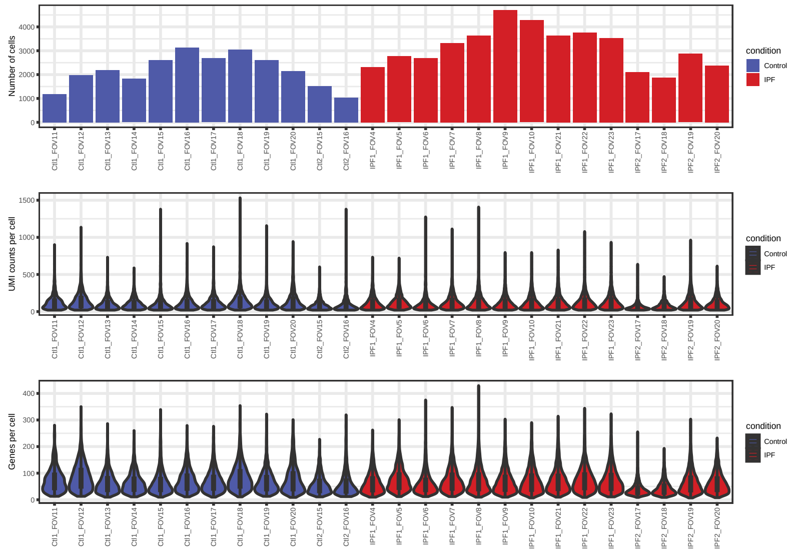

### sup2

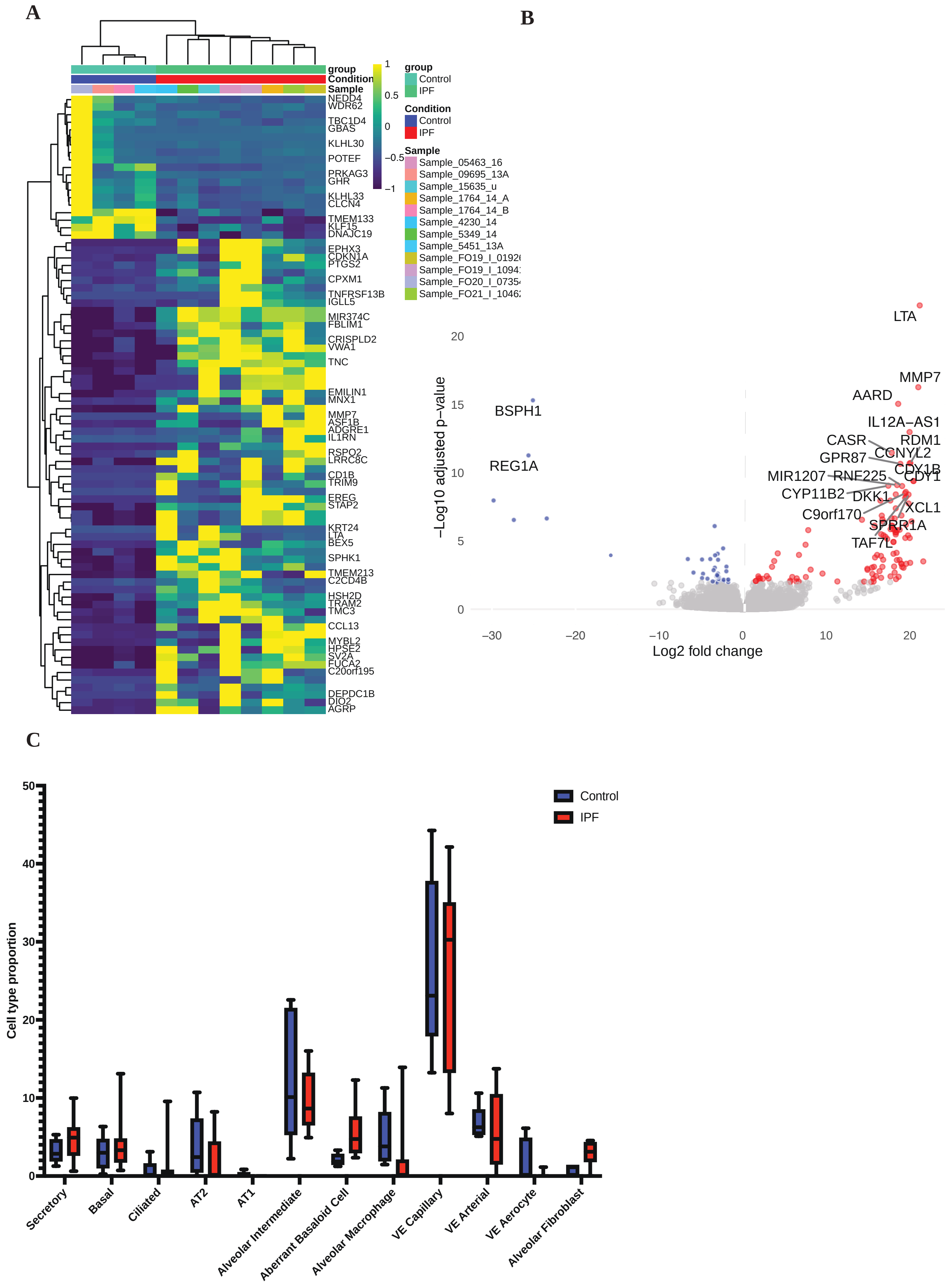

### sup4

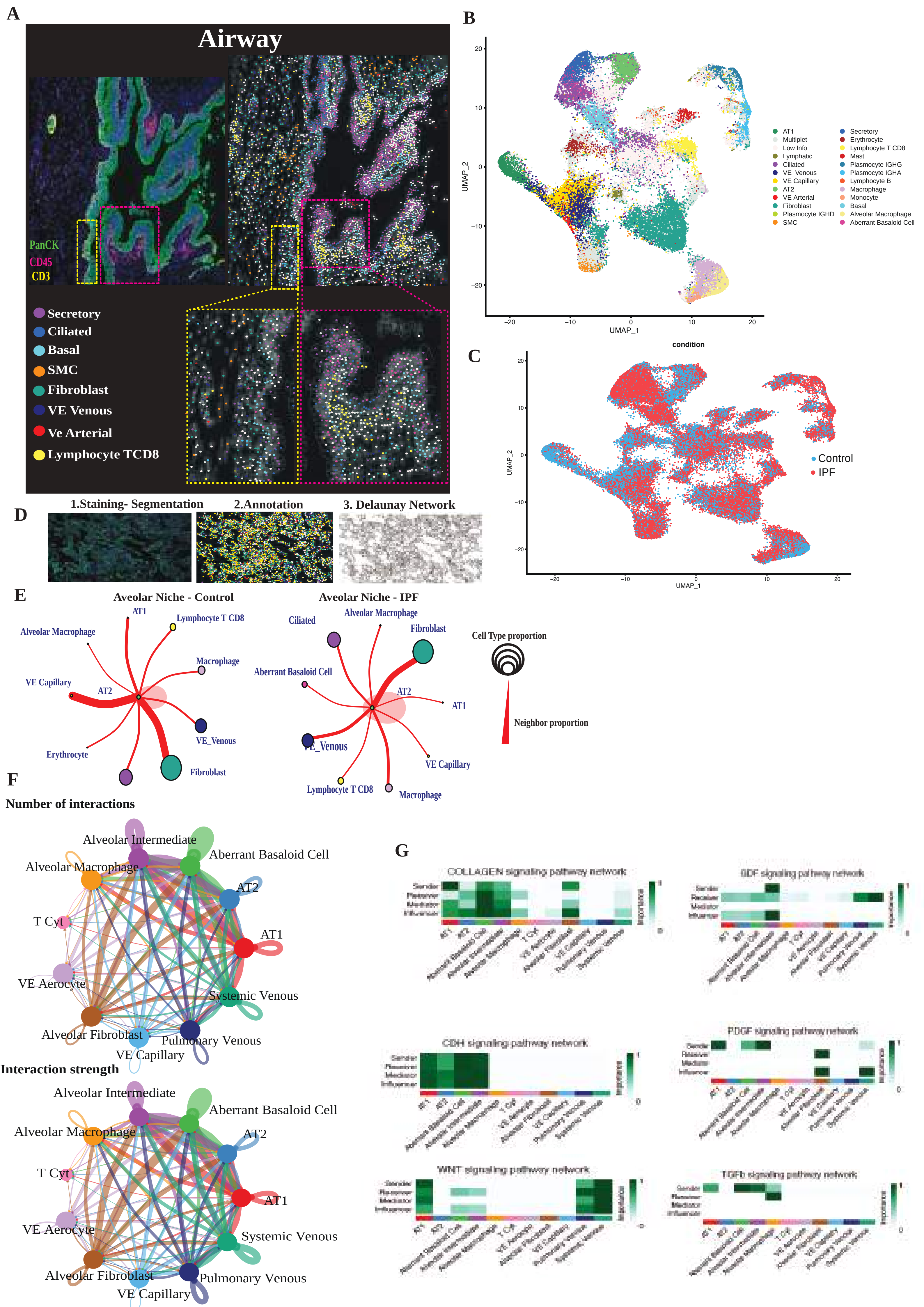
