## Supplementary material for "Alveolar niche disruption and aberrant epithelial reprogramming are early hallmarks of idiopathic pulmonary fibrosis": sup3

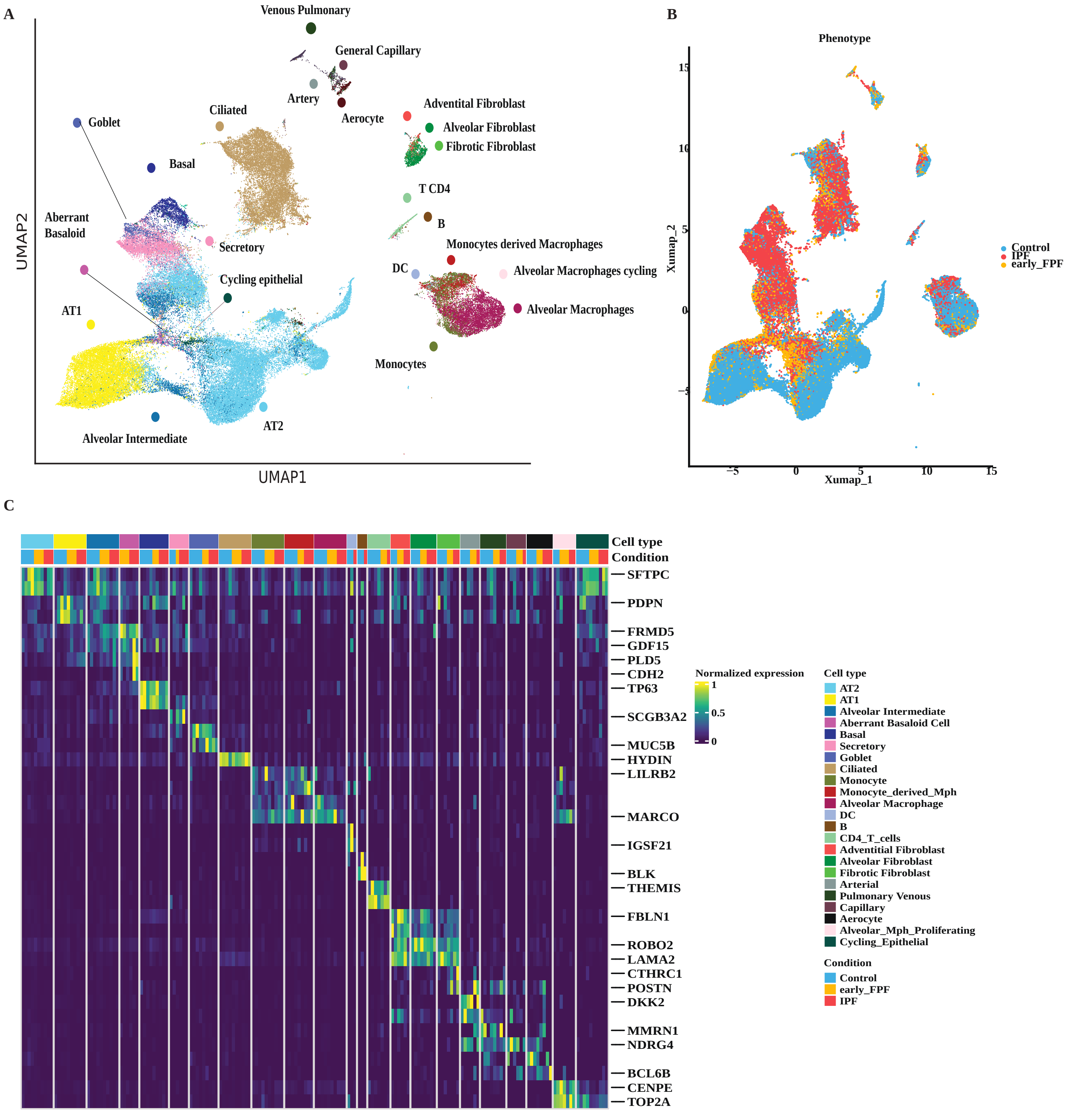


**D**

| | AT1 | AT2 | Aberrant Basaloid Cell | Adventitial Fibroblast | Alveolar Fibroblast | Alveolar Intermediate | Alveolar Macrophage | B | Basal | Ciliated | DC | Aerocyte | Arterial | Capillary | Pulmonary Venous | Fibrotic Fibroblast | Goblet | Lymphatic | Mast | Monocyte | Secretory |
|---|---|---|---|---|---|---|---|---|---|---|---|---|---|---|---|---|---|---|---|---|---|
| AT1 | 0.97 | 0.09 | 0.25 | -0.13 | -0.13 | 0.37 | -0.06 | -0.13 | -0.03 | 0.00 | -0.16 | -0.02 | -0.12 | -0.07 | -0.02 | -0.19 | -0.06 | -0.05 | -0.09 | -0.13 | -0.07 |
| AT2 | 0.08 | 0.97 | 0.21 | -0.15 | -0.10 | 0.60 | -0.06 | -0.12 | -0.03 | -0.01 | -0.11 | -0.07 | -0.13 | -0.05 | 0.05 | -0.21 | -0.03 | -0.06 | -0.16 | -0.16 | -0.02 |
| Aberrant Basaloid Cell | 0.17 | 0.17 | 0.86 | -0.15 | -0.10 | 0.37 | -0.13 | 0.01 | 0.27 | -0.03 | -0.15 | -0.15 | -0.20 | -0.24 | -0.12 | -0.09 | -0.09 | -0.11 | -0.09 | -0.14 | -0.04 |
| Adventitial Fibroblast | -0.12 | -0.17 | -0.18 | 0.88 | 0.51 | -0.20 | -0.08 | -0.12 | -0.12 | -0.09 | -0.10 | -0.17 | -0.14 | -0.22 | -0.17 | 0.55 | -0.10 | 0.04 | -0.11 | -0.04 | -0.13 |
| Alveolar Fibroblast | -0.11 | -0.13 | -0.19 | 0.49 | 0.90 | -0.19 | 0.01 | -0.14 | -0.14 | -0.08 | -0.13 | -0.13 | -0.09 | -0.17 | -0.19 | 0.63 | -0.13 | -0.11 | -0.06 | -0.05 | -0.15 |
| Alveolar Intermediate | 0.47 | 0.42 | 0.77 | -0.21 | -0.18 | 0.74 | -0.13 | -0.11 | 0.10 | -0.01 | -0.17 | -0.13 | -0.23 | -0.20 | -0.08 | -0.23 | -0.07 | -0.09 | -0.13 | -0.21 | -0.04 |
| Alveolar Macrophage | -0.06 | -0.05 | -0.15 | -0.11 | -0.09 | -0.12 | 0.92 | 0.10 | -0.11 | -0.03 | 0.13 | -0.08 | -0.11 | -0.07 | -0.14 | -0.13 | -0.08 | -0.10 | -0.02 | 0.70 | -0.08 |
| B | -0.08 | -0.09 | -0.17 | -0.11 | -0.08 | -0.18 | 0.06 | 0.73 | -0.14 | -0.01 | 0.22 | 0.00 | -0.08 | -0.09 | -0.13 | -0.13 | -0.13 | -0.09 | 0.26 | 0.13 | -0.13 |
| Basal | -0.04 | 0.01 | 0.14 | -0.14 | -0.06 | 0.11 | -0.09 | -0.11 | 0.91 | 0.00 | -0.05 | -0.14 | -0.13 | -0.19 | -0.17 | -0.02 | 0.13 | -0.13 | -0.06 | -0.10 | 0.17 |
| Ciliated | -0.04 | 0.01 | -0.03 | -0.09 | -0.12 | 0.01 | -0.07 | -0.04 | 0.12 | 0.96 | 0.00 | -0.10 | -0.12 | -0.14 | -0.16 | -0.09 | 0.08 | -0.07 | -0.09 | -0.04 | 0.09 |
| DC | -0.15 | -0.13 | -0.22 | -0.12 | -0.09 | -0.25 | 0.19 | 0.43 | -0.15 | -0.06 | 0.81 | -0.06 | -0.10 | -0.03 | -0.12 | -0.11 | -0.16 | -0.10 | 0.21 | 0.36 | -0.16 |
| Aerocyte | -0.10 | -0.11 | -0.17 | -0.14 | -0.14 | -0.19 | -0.05 | -0.01 | -0.15 | -0.07 | -0.05 | 0.93 | 0.09 | 0.46 | 0.28 | -0.15 | -0.12 | 0.07 | -0.10 | -0.06 | -0.14 |
| Arterial | -0.19 | -0.22 | -0.28 | 0.11 | 0.21 | -0.29 | -0.08 | -0.18 | -0.24 | -0.15 | -0.17 | 0.14 | 0.79 | 0.49 | 0.33 | 0.24 | -0.18 | -0.04 | -0.09 | -0.11 | -0.19 |
| Capillary | -0.13 | -0.14 | -0.19 | -0.21 | -0.22 | -0.20 | -0.07 | -0.14 | -0.18 | -0.11 | -0.12 | 0.51 | 0.51 | 0.92 | 0.51 | -0.19 | -0.11 | 0.03 | -0.14 | -0.11 | -0.15 |
| Pulmonary Venous | -0.16 | -0.18 | -0.22 | -0.14 | -0.18 | -0.26 | -0.13 | -0.19 | -0.20 | -0.14 | -0.15 | 0.28 | 0.61 | 0.67 | 0.88 | -0.15 | -0.10 | 0.20 | -0.12 | -0.15 | -0.14 |
| Fibrotic Fibroblast | -0.17 | -0.20 | -0.18 | 0.51 | 0.59 | -0.25 | -0.06 | -0.08 | -0.14 | -0.10 | -0.05 | -0.16 | -0.02 | -0.22 | -0.14 | 0.79 | -0.13 | -0.09 | -0.01 | -0.01 | -0.10 |
| Goblet | -0.09 | -0.03 | -0.09 | -0.19 | -0.22 | -0.04 | -0.10 | -0.10 | 0.20 | 0.01 | -0.05 | -0.17 | -0.06 | -0.18 | -0.15 | -0.18 | 0.94 | -0.12 | -0.12 | -0.11 | 0.87 |
| Lymphatic | -0.09 | -0.11 | -0.13 | -0.03 | -0.12 | -0.14 | -0.06 | -0.09 | -0.11 | -0.07 | -0.09 | 0.00 | 0.03 | -0.02 | 0.16 | -0.08 | -0.09 | 0.96 | -0.04 | -0.06 | -0.10 |
| Mast | -0.05 | -0.06 | -0.13 | -0.09 | -0.12 | -0.12 | 0.00 | 0.24 | -0.11 | -0.03 | 0.13 | -0.06 | -0.09 | -0.11 | -0.15 | -0.12 | -0.11 | -0.13 | 0.80 | 0.05 | -0.10 |
| Monocyte | -0.08 | -0.08 | -0.18 | -0.11 | -0.11 | -0.20 | 0.23 | 0.24 | -0.16 | -0.01 | 0.66 | -0.08 | -0.13 | -0.07 | -0.16 | -0.12 | -0.10 | -0.13 | 0.22 | 0.66 | -0.12 |
| Secretory | 0.08 | 0.17 | 0.33 | -0.16 | -0.22 | 0.64 | -0.08 | -0.15 | 0.34 | 0.05 | -0.09 | -0.22 | -0.18 | -0.26 | -0.23 | -0.19 | 0.34 | -0.10 | -0.15 | -0.17 | 0.50 |
